# Measuring graded changes in consciousness through multi-target filling-in

**DOI:** 10.1101/499517

**Authors:** Matthew J Davidson, Irene Graafsma, Naotsugu Tsuchiya, Jeroen van Boxtel

## Abstract

Perceptual filling-in (PFI) occurs when a physically-present visual target disappears from conscious perception, with its location filled in by the surrounding visual background. Compared to other visual illusions, these perceptual changes are crisp and simple, and can occur for multiple spatially-separated targets simultaneously. Contrasting neural activity during the presence or absence of PFI may complement other multistable phenomena to reveal the neural correlates of consciousness (NCC). We presented four peripheral targets over a background dynamically updating at 20 Hz. While participants reported on target disappearances/reappearances via button press/release, we tracked neural activity entrained by the background during PFI using steady-state visually evoked potentials (SSVEPs) recorded in the electroencephalogram. We found background SSVEPs closely correlated with subjective report, and increased with an increasing amount of PFI. Unexpectedly, we found that as the number of filled-in targets increased, the duration of target disappearances also increased, suggesting facilitatory interactions exist between targets in separate visual quadrants. We also found distinct spatiotemporal correlates for the background SSVEP harmonics. Prior to genuine PFI, the response at the second harmonic (40 Hz) increased before the first (20 Hz), which we tentatively link to an attentional effect. There was no difference between harmonics for physically removed stimuli. These results demonstrate that PFI can be used to study multi-object perceptual suppression when frequency-tagging the background of a visual display, and because there are distinct neural correlates for endogenously and exogenously induced changes in consciousness, that it is ideally suited to study the NCC.

**Highlights:**

- Perceptual filling-in (PFI) has distinct advantages for investigating the neural correlates of consciousness.
- Participants can accurately report graded changes in consciousness using four simultaneous buttons.
- Frequency-tagging of visual background information tracks changes in visual perception.
- Spatiotemporal EEG responses differentiate PFI from phenomenally matched physical disappearances.

## Introduction

In perceptual filling-in (PFI) phenomena, areas of the visual environment that are physically distinct become interpolated by the visual features of the surrounding texture or background (Komatsu, 2006; Meng, Remus, & Tong, 2005; Pessoa, Thompson, & Noë, 1998; Ramachandran & Gregory, 1991; Weil & Rees, 2011). Although PFI neatly displays how our awareness of a visual scene is shaped by unconscious inferential processes (Komatsu, 2006), it has traditionally been investigated to understand how our visual system compensates for retinal-blind spots (Durgin, Tripathy, & Levi, 1995; Komatsu, Kinoshita, & Murakami, 2000; Ramachandran & Gregory, 1991; Spillmann, Otte, Hamburger, & Magnussen, 2006), and visual field defects (Gassel & Williams, 1963; Gerrits & Timmerman, 1969; Safran & Landis, 1996). Accordingly, a range of low-level visual attributes such as target contrast (Stürzel & Spillmann, 2001) target eccentricity (De Weerd, Desimone, & Ungerleider, 1998), and microsaccades (Martinez-Conde, Macknik, Troncoso, & Dyar, 2006; Troncoso, Macknik, & Martinez-Conde, 2008) have been shown to affect the dynamics of PFI. As a result, the neural interpolation of information in lower visual areas has been implicated as one active mechanism behind PFI (De Weerd, Gattass, Desimone, & Ungerleider, 1995; Komatsu, 2006; Meng et al., 2005; Pessoa et al., 1998).

In addition to the role of low-level visual processes, top-down attention and higher-cortical areas have also been implied to play a role in the initiation, maintenance, and termination of PFI (De Weerd, Smith, & Greenberg, 2006; Weil, Wykes, Carmel, & Rees, 2012). For example, selectively attending to the location of a target (De Weerd, 2006; De Weerd et al., 2006), or attending to shared features among peripheral targets (De Weerd et al., 2006; Lou, 1999) has been shown to increase the likelihood of PFI. This poses an intriguing puzzle, as neural responses to a sensory stimulus usually increase when prioritized by top-down attention (Harris & Thiele, 2011; Reynolds, Pasternak, & Desimone, 2000; Spitzer, Desimone, & Moran, 1988) and also increase when the stimulus is consciously perceived (e.g. De Weerd et al., 1995; Polonsky, Blake, Braun, & Heeger, 2000 but also see Donner, Sagi, Bonneh, & Heeger, 2008; Logothetis, 1998; Watanabe et al., 2011). As attention during PFI increases disappearance rates, insights into the mechanisms of this phenomenon may contribute to the hotly debated dissociation between attention and consciousness (Koch & Tsuchiya, 2007; Ling & Carrasco, 2006; van Boxtel, Tsuchiya, & Koch, 2010a, 2010b). We were also motivated to explore whether the use of PFI could capture graded changes to the neural correlates of conscious perception (Windey & Cleeremans, 2015; Windey, Vermeiren, Atas, & Cleeremans, 2014), as PFI can occur over multiple regions embedded in the same visual-background. More specifically, we hypothesized that upon the phenomenological interpolation of target regions, an increased neural response to the surrounding visual background would be recorded, consistent with active mechanism accounts of perceptual suppression during PFI (De Weerd et al., 1995; Komatsu, 2006; Meng et al., 2005; Pessoa et al., 1998).

We investigated the neural correlates of PFI through the use of frequency-tagging in the EEG, and focused on the background of our visual display in contrast to a previous study that looked only at a single target stimulus (Weil, Kilner, Haynes, & Rees, 2007). By presenting flickering visual stimuli, frequency-tagging elicits a steady-state visually evoked potential (SSVEP), which can be analysed as a narrowband change in power at the flicker-frequency of interest (Norcia, Appelbaum, Ales, Cottereau, & Rossion, 2015; Vialatte, Maurice, Dauwels, & Cichocki, 2010). This flicker effect is used to ‘tag’ populations of neurons processing each flickering stimulus (reviewed in Norcia et al., 2015). SSVEPs have been used to track fluctuations in visual awareness between competing stimuli (Brown & Norcia, 1997; Katyal, Engel, He, & He, 2016; Lansing, 1964; Sutoyo & Srinivasan, 2009; Tononi, Srinivasan, Russell, & Edelman, 1998; Zhang, Jamison, Engel, He, & He, 2011) as well as to track the allocation of attention (Andersen, Hillyard, & Müller, 2008; Müller et al., 2006; Müller, Teder-Salejarvi, & Hillyard, 1998; Müller & Hillyard, 2000; Müller & Hübner, 2002). The latter effect may be particularly strong in the second harmonic (i.e. frequency double) of the SSVEP driving frequency (Kim, Grabowecky, Paller, Muthu, & Suzuki, 2007; Kim, Grabowecky, Paller, & Suzuki, 2011; Pei, Pettet, & Norcia, 2002). To investigate the neural correlates of PFI, we combined the SSVEP technique with a novel multi-target PFI paradigm, by frequency-tagging the background of our visual display. This approach allowed us to obtain a more graded response to the amount of change in conscious perception, by investigating how the number of targets perceptually filled-in influenced neural responses to the shared visual-background display.

## Methods and Materials

### Participants

Twenty-nine healthy volunteers (11 male, 18-39 years of age, *M* = 24, *SD* = 5 years) took part in the study. As the effect size of frequency-tagging the background-stimuli is unknown, we chose to test twice as many participants than previous studies that used frequency-tagging to study the neural correlates of perception during binocular rivalry (e.g. Katyal et al., 2016; Zhang et al., 2011), as well as target-responses during PFI (Weil et al., 2007). Participants had normal or corrected-to-normal vision. All participants were recruited via convenience sampling, provided written informed consent prior to participation, and received a monetary compensation (30 AUD) for their time. The study was approved by the Monash University Human Research and Ethics Committee (MUHREC #CLF016).

## Apparatus and stimuli

Participants were seated in a dark room approximately 50 cm distance from a computer monitor (size 29 x 51 cm, resolution 1080 x 1920 pixels, subtending 32 x 54° visual angle, refresh rate 60 Hz). To frequency-tag the background image, we prepared 100 random patterns prior to the start of each experiment. To construct each background pattern, we first down-sampled the screen to 540 x 960 pixels. Then we assigned a random luminance value (drawn from a uniform distribution from black to white) to each down-sampled pixel. These background images were refreshed at a rate of 20 Hz by randomly selecting from the set of 100 prepared patterns.

On top of this background image, the display was composed of a central fixation cross (1.03° visual angle in height and width), surrounded by four counter-phase flickering 2 x 2 checkerboard targets (4.56° visual angle in diameter). A target was located in each quadrant, centred at 23.3° eccentricity from the centre of the screen. The targets were located closer to the horizontal than vertical (horizontal distance from centre 20.2°, vertical distance from centre 11.5°; Figure 1). Targets were smoothly alpha-blended into the background texture following a 2D Gaussian profile (*SD* = 1.06° visual angle in diameter). As small, peripherally located targets flickering above 7 Hz are more likely to disappear (Anstis, 1996; Schieting & Spillmann, 1987) each target was also flickered by reversing the contrast of checkerboard elements at one of four unique frequencies (8, 13, 15 and 18 Hz).

**Figure 1:**
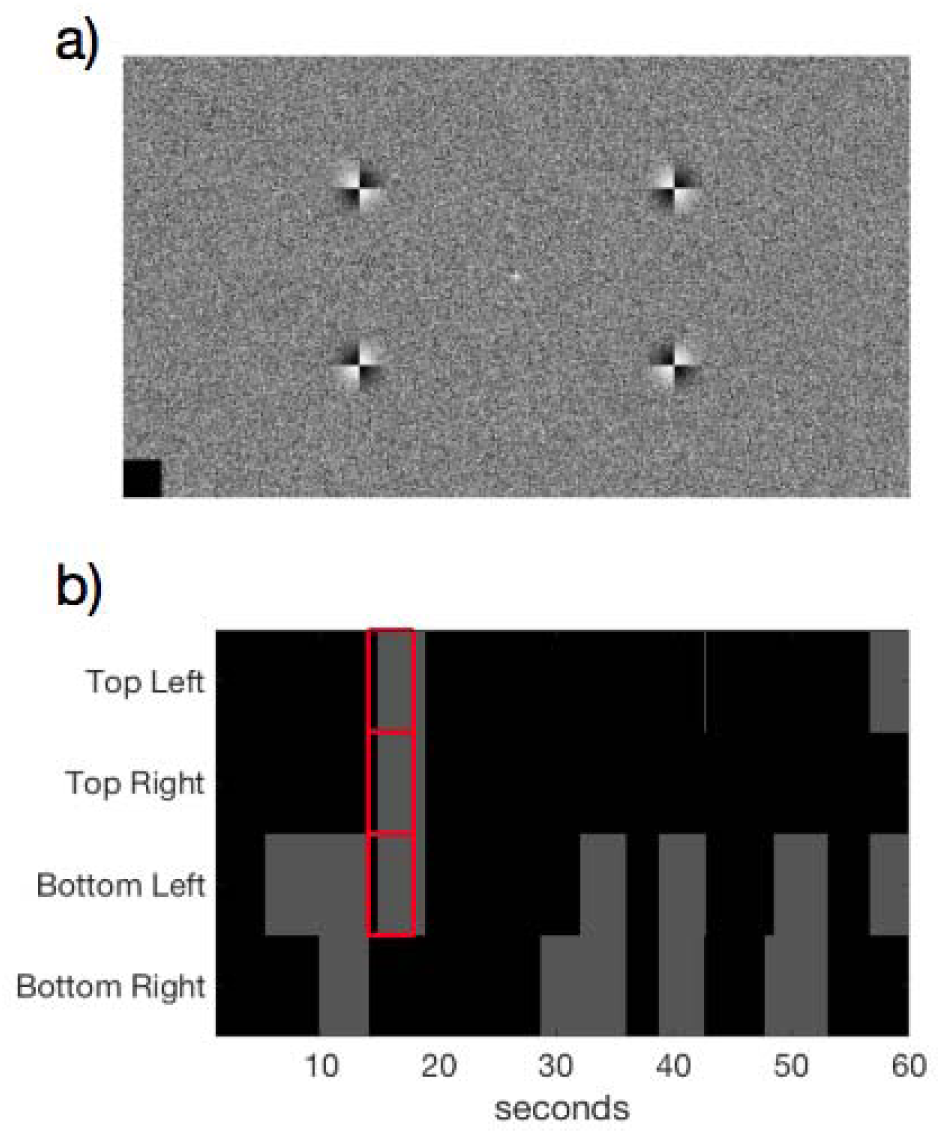
Stimulus display and example response. a) Stimulus display containing a central fixation cross, dynamic background (updated at 20 Hz) and four target checkerboard stimuli. b) Example time course of behavioural responses over a 60-second trial from one participant. Participants were asked to monitor each peripheral target simultaneously, and to press and hold each button upon perceptual disappearance (PFI events shown in grey) at the corresponding location of the target. PMD periods are shown in red, for which targets were physically replaced by the flickering background texture. Note that targets often disappear and reappear together. An example trial of this display can be viewed at https://figshare.com/articles/Example_trial/7320233.

We chose to use small, peripherally located flickering targets to optimally induce disappearances across all our participants. Yet, these same parameters are sub-optimal for frequency-tagging stimuli. As a result, our simultaneous background flicker remains as a relatively pure EEG measure for tracking the disappearance of multiple disappearing PFI targets, which is ideal because the measure of background SSVEP is a novel element of our experiment.

### Task procedure

Each experimental session was composed of 25 trials, 60 seconds per trial. Between the trials, participants were able to take short self-timed breaks, resulting in a total time-on-task of approximately 30 minutes. Before the experiment, participants were instructed to fixate on the central cross, and were informed that they may sometimes experience a visual illusion where any number of peripheral targets may disappear from their field of vision. We did not monitor eye position. Participants then completed one practice trial to familiarize themselves with the corresponding button presses required for targets in each of the four visual quadrants. Specifically, they were instructed to press keys ‘A’, ‘Z’, ‘K’, and ‘M’ on a traditional QWERTY keyboard, assigning them to the upper left, bottom left, upper right, and bottom right targets, respectively. Participants were instructed to hold each button for the duration of disappearance of the corresponding target, and to release it immediately upon the corresponding reappearance. Figure 1 presents the basic configuration of the experimental display used (see Movie 1 for an example of the flickering background display).

### Phenomenally matched disappearance (PMD) periods

We introduced PMD periods to check if participants were correctly reporting on disappearances. During a PMD period 1 to 4 targets were physically removed from the display and replaced with the background through alpha blending. Multiple targets during PMD periods were removed with the same onset. Each PMD period lasted from 3.5 to 5 seconds in duration (drawn from a uniform distribution). To mimic the phenomenology of endogenous PFI events, we generated PMD periods by linearly ramping the luminance contrast of the target up or down over 1.5 seconds. Participants were not informed of the PMD periods.

These physical PMD periods also served as a control condition for comparison with the neural signals evoked by PFI. Within 25 trials, PMD events in which one, two, three or four targets were removed each occurred on six trials for each participant. The location of the removed targets in the case of one, two and three targets were randomized. The order of these PMD events were also randomized for each experiment. A previous study showed that flickering peripheral targets tend not to disappear in the beginning of trials (Schieting & Spillmann, 1987), so each PMD event began no sooner than 10 seconds after the beginning of each trial to ensure that PMD disappearances remained indistinguishable from PFI. Our own data also confirmed that participants reported much lower PFI in the initial 10 seconds of each trial, with PFI plateauing after approximately 10-15 seconds. We also did not include PMDs within the last 10 seconds. We note that for 10 of our 29 participants, four-target PMD periods did not occur due to a coding error, and instead all four targets remained on screen, resulting in PMD periods being presented on 92% of trials overall (over all *N*=29 participants).

### Participant and trial exclusion based on PMD periods

Initial screening analyses sought to confirm whether participants were able to simultaneously monitor the visibility of multiple peripheral targets using four unique buttons, and perform this task accurately and in compliance with instructions. Due to a keyboard malfunction, button press responses to three and four disappearing targets became indistinguishable in our post-hoc analysis, and have been analysed together henceforth as “3 or 4 buttons pressed”. In the subsequent analyses where the number of buttons pressed mattered, we proceeded as if three buttons were pressed in these periods.

We analysed the likelihood of button press responses during PMD periods to estimate participant attention on task. As PMD periods were embedded within a trial, some PMD periods occurred when participants had already pressed buttons. Such events are more frequent for those who report more frequent PFI, and accounting for this baseline likelihood of button press is necessary to accurately estimate PMD-responses. To estimate this baseline button press rate per individual participant, we performed a bootstrapping analysis with replacement. For a given PMD onset in trial T at time S (seconds), we randomly selected a trial T’ (T=T’ was allowed) and epoched the button press time course over the period of [S-2, S+4] at corresponding PMD target locations in trial T’. We repeated this for all trials (T=1…24, except for the 4-PMD error mentioned above) to obtain a single bootstrapped set of trials per participant. We then obtained the mean button press time course across button-locations from this bootstrapped set, and repeated the procedure 200 times to obtain a null distribution, representing the likelihood of baseline button press around PMD onset per participant (Figure 2a, grey lines and shading). We also obtained the mean button press time course for observed data across button-locations.

**Figure 2.**
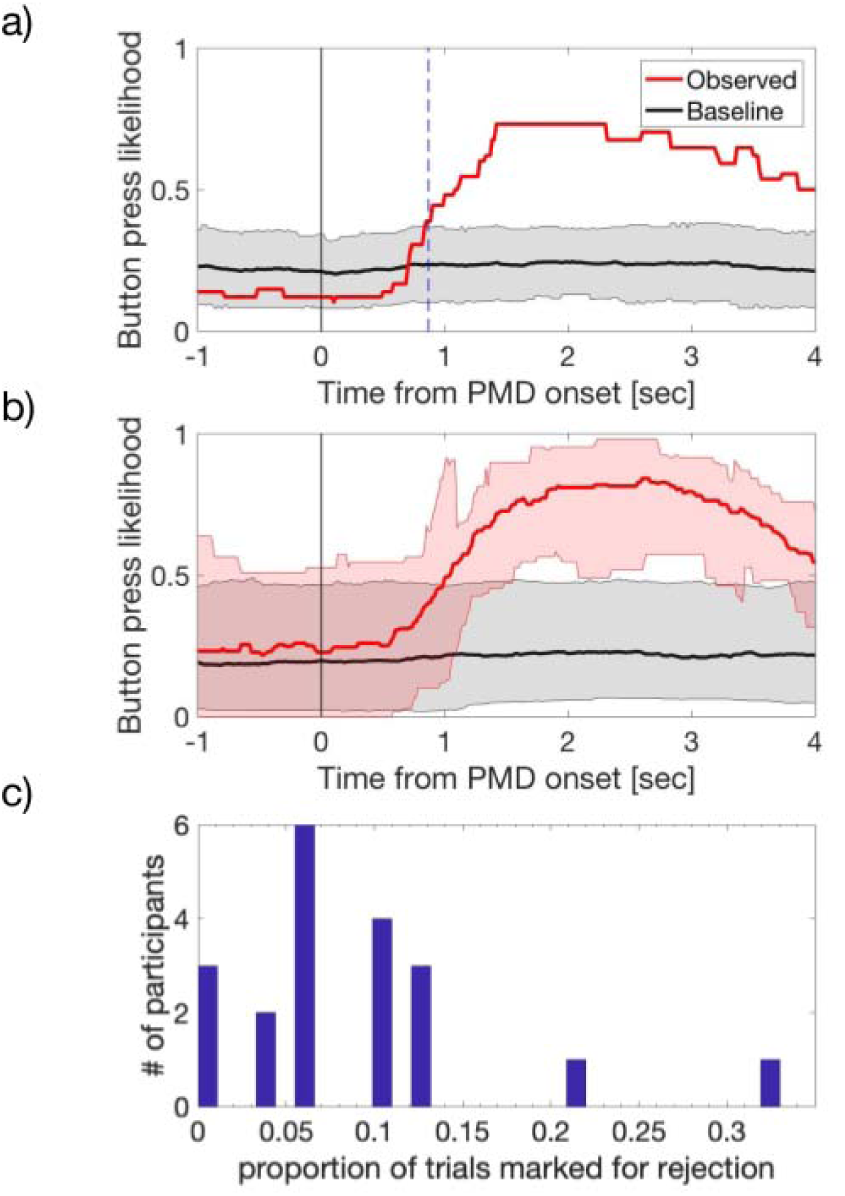
PMD period analysis and trial rejection following the physical removal of flickering targets at PMD onset. a-b) display the likelihood of button press time-courses for observed (red) and bootstrapped data (grey). a) Example PMD response for a single participant. The first time point that the observed likelihood of button press (red) exceeded the bootstrapped CI (grey) corresponds to the PMD reaction time (0.87 second for this participant, marked with a vertical dashed blue line). b) The mean time course for the likelihood of button press and its bootstrapped sets across participants, shown in red and grey respectively. Shading represent the CI (computed with logit transform and presented after reverse transform) across participants. c) Participant-level histogram of the proportion of trials rejected, based on period-by-period PMD analysis.

As the distribution at each time point for both observed and baseline data was not normally distributed, we first converted the data into z-scores using the logit transformation before calculating the confidence intervals (CI). Then, we used mean z-scores (±1.96 *SD* of z-scores) as the CI for the null distribution of baseline data within each participant, and observed data across participants.

We excluded three participants whose mean button press time course around the actual PMD onset failed to exceed the CI of the baseline button press likelihood within the first two seconds (i.e., [0, S+2]). We defined the PMD onset reaction time as the first time point after which the mean button press data exceeded the top CI, indicating successful button presses for PMD targets. Figure 2a shows the PMD response for an example participant retained for analysis. Four further participants were removed from subsequent analyses for failing to experience PFI during most of the experimental session (i.e., only brief events on 1 or 2 trials). For the remaining participants, the mean reaction time to respond to PMD onsets, and thus the disappearance of a peripheral target was 0.92 seconds (*SD* = 0.046). Figure 2b shows the proportion of button press responses for all PMD events across participants retained for analysis (*N*=22).

Having identified which participants could successfully indicate target disappearance based on their button press data, we continued to identify and remove any trials from the subsequent analysis in which a PMD was not correctly detected. We undertook this procedure to assure that in all retained trials participants paid proper attention on task and reported accurately on PFI. We regarded a PMD period as being successfully identified if participants pressed the corresponding button for at least 50% of the allowed response time window. For multi-target PMD periods, we applied the same criteria for each button separately. If any button was not pressed at least 50% of the time, the PMD was considered undetected. For four-target PMD periods, we analysed it as if it was a three-target PMD period. This window was from the onset of the PMD plus 1 second (in consideration of the reaction time delay) to the end of PMD. For example, if the PMD period under consideration was 3.5 seconds in duration, we defined the allowed time window to be [1, 3.5] seconds from the PMD onset. Figure 2c shows a participant-level histogram for the number of rejected trials (*M ± SD:* 1.75 *±* 1.48 trials or 8.96 *±* 7.89% of all trials). After participant and trial exclusion, we continued by examining the behavioural dynamics of PFI.

### Quantifying PFI and PFI location-shuffling analysis

We analysed the number of PFI events, duration of each PFI event, and total duration of PFI per 60 second trial. Although these variables may be correlated, they have also been shown to reveal complementary data features in similar multi-target designs e.g. (Bonneh, Donner, Cooperman, Heeger, & Sagi, 2014; McEwen, Paton, Tsuchiya, & van Boxtel, 2018; Thomas, Davidson, Zakavi, Tsuchiya, & van Boxtel, 2017). Each of these variables were compared based on the number of PFI (nPFI; 1, 2, 3 or 4), quantified by the number of simultaneous buttons pressed at each time point. We note that our reported analysis did not exclude transient button press periods (<200 ms), as the dynamics of multi-target PFI are currently unknown. Instead, we performed a reconstruction analysis to estimate the contribution of overlapping and closely spaced PFI events on EEG responses (see ***Reconstruction analysis - Methods).*** However, when we repeated our behavioural analysis by removing these instances (< 200 ms), the results were not significantly different.

To investigate whether the simultaneous multi-target PFI observed in participant data (e.g. Figure 1b) exceeded that expected by chance, we performed a shuffling analysis to create a null distribution. Specifically, we created 1000 shuffled trials for each participant, by randomly selecting the button press time course for each of the four target locations independently from any of the trials throughout their experimental session (this could include multiple locations within the same trial). As such, newly created shuffled trials allowed us to compare the effect of multiple disappearing target events within the same trial (the observed data) to the shuffled data without the presence of a temporal correlation between target locations. If target disappearances during PFI were independent, then shuffled and experimental data should be similar. The comparison between the observed and the shuffled data is displayed in Figure 7.

### Linear-mixed effect analysis – Behaviour

All statistical analyses were performed using MATLAB (ver: R2016b). We used linear-mixed effect (LME) analysis to examine whether various PFI characteristics (e.g., durations) were affected by the number of simultaneously invisible targets (nPFI; *n* = 0, 1, 2, 3 or 4), including intercepts for participants as a random effect. We performed likelihood ratio tests between the full model and a restricted model which excluded the factor of interest (Glover & Dixon, 2004; Pinheiro, Bates, DebRoy, & Sarkar, 2014; Winter, 2013a, 2013b).

We also performed LME analyses to compare the slopes of observed and shuffled data, when considering the effect of the number of simultaneously invisible targets on PFI characteristics. For this analysis, we fit a linear model (1st order polynomial) to the observed data across participants (*N* = 22), and retained the slope (β) as our observed test statistic. Similarly, we also fit the same linear model to each of *n* = 1000 sets of shuffled data, each of which was computed from the shuffled trials across *N* = 22 participants. We shuffled the trials within each participant within each set and again retained the β values. Then, we compared the observed β value with the null distribution of the β values from *n* = 1000 shuffled sets. If the observed β exceeded the top 97.5% or was lower than 2.5% of the null distribution, we considered the observed effect to be significant at *p* < .05.

### EEG acquisition and pre-processing

Throughout each session whole-head EEG was recorded with 64 active electrodes arranged across an elastic-cap according to the international 10-10 system. Electrode impedances were kept below 10 kΩ prior to experimentation, and recorded using the default reference (FCz) and ground electrode (AFz) via Brainvision recorder software (sampling rate = 1000 Hz, offline bandpass of 0.5-70 Hz). All EEG data was stored for offline analysis using custom MATLAB scripts (Ver: R2016b), as well as the EEGLAB (Delorme & Makeig, 2004) and Chronux (Bokil, Andrews, Kulkarni, Mehta, & Mitra, 2010) toolboxes. All EEG channels were first re-referenced to the average of all electrodes at each sample and down-sampled to 250 Hz. We further applied a Laplacian transform to improve spatial selectivity of the data, which is known to contribute minimal contamination to the SSVEP when using rhythmic-entrainment source separation (RESS; Cohen & Gulbinaite, 2017), which we used to extract SSVEP responses as detailed below.

### SSVEP Signal-to-Noise Ratio (SNR) calculation

To estimate the topography and across channel correlation of SSVEPs (Figures 5 and 11), we first calculated the natural log of the power spectrum via the fast Fourier transform (FFT) over the period −3000 to −100 ms before, and 100 to 3000 ms after button press. We excluded the 200 ms around button press responses to avoid motor-evoked activity. In the SSVEP paradigm, we operationally regard power at the tagged frequency as signal and power at non-tagged neighbouring frequencies as noise (Norcia et al., 2015), and compute the signal-to-noise ratio (SNR) at each frequency. In logarithmic scale, this corresponds to log of the power at each frequency subtracted by the mean log power across the neighbourhood frequencies. In this paper, all SNR results are based on this log-transformed SNR metric because without log-transform, SNR is highly skewed and not appropriate for various statistical tests. Over the 2.9 s time window (half-bandwidth = 0.35Hz), we computed the SNR at frequency f (Hz) as the mean log power over the neighbourhood frequencies for f subtracted from the log power at f. Neighbourhood frequencies began at least one-half bandwidth from f, to avoid capturing stimulus related signal in the SNR calculation. The neighbourhood for power spectra defined above was defined as [f-1.22, f-0.44] Hz and [f+0.44, f+1.22] Hz. In addition, we also computed the time-course of the SNR over a 1 second window (half-bandwidth = 1 Hz) with a step-size of 0.15 second, to enable the comparison of fluctuations in SNR over time. For this shorter time window, we used the neighbourhood as [f-3.92, f-1.95] Hz and [f+1.95, f+3.92] Hz to compute the log(SNR) time course.

### SSVEP analysis via rhythmic entrainment source separation (RESS)

After examining the topography of log(SNR) responses, we applied rhythmic entrainment source separation (RESS) to optimally extract the time-course of frequency-tagged components of SSVEPs without relying upon electrode channel selection (Cohen & Gulbinaite, 2016). In standard SSVEP analysis, the SNR is examined by averaging across electrodes within a region of interest or selecting one electrode in a certain way (e.g., prior hypothesis, anatomical localization or separate datasets). An alternative to this classic approach is RESS, which creates a map of spatial weights across all electrodes which optimize the SNR at a particular frequency, tailored for each participant. Specifically, RESS functions by creating linear spatial filters to maximally differentiate the covariance between a signal flicker frequency and neighbourhood frequencies, thereby increasing the signal-to-noise ratio at the flicker frequency. After obtaining signal and neighbourhood covariance matrices, the eigenvector with the largest eigenvalue is used as channel weights to reduce the dimensionality of multi-channel data into a single component time course, which reduces multiple comparisons across channels in statistical testing.

After epoching all data using the time-windows −3000 to −100 ms and 100 ms to 3000 ms peri button press/release, we then constructed RESS spatial filters per participant, avoiding PMD periods. We constructed RESS spatial filters from 64-channel EEG, by extracting signal data following a narrow-band filter via frequency-domain Gaussian, centred at flicker frequencies (20 and 40 Hz, full-width at half maximum = 1 Hz). We proceeded by selecting broadband (unfiltered) neural activity to construct reference covariance matrices. We selected broadband EEG as our reference after confirming that improvements to the SNR at both the 20 and 40 Hz signal were statistically equivalent. Comparing signal to broadband activity as a reference, as opposed to immediate neighbourhood frequencies, has previously been shown to allow the optimization of SSVEP signals using RESS (Cohen & Gulbinaite, 2017).

Critically, we performed the above procedure without distinguishing whether targets were disappearing or reappearing due to button press or release in order to reduce the possibility of overfitting. If we were to construct separate filters for periods around the time of target disappearance and reappearance, then any differences between these conditions could be due to differences in the obtained filters, or overfitting of the filters prior to our condition comparisons. After application of the RESS spatial filters, we calculated the time course of SSVEP log(SNR) from the RESS component time courses, separately for each background flicker of interest as described above (***SSVEP Signal-to-Noise Ratio (SNR) calculation***). The main results of our analysis also hold with more conventional SSVEP analysis, such as when focusing on only parieto-occipital electrodes.

### SNR time-course data cleaning

Preliminary analyses revealed a sharp and consistent decrease in 40 Hz log(SNR) amplitude which was time-locked to the beginning of each PMD period. Subsequent inspection of recorded screen flip-times revealed a lag in background stimulus presentation (16.7-33.3 ms duration) at PMD onset, which resulted in the background pixels for one presentation frame being skipped. This caused an artefact in the spectrogram where the time window of the analysis included the problematic period. To correct for this artefact conservatively, we interpolated the 40 Hz SNR time-course from −500 to 500 ms around physical PMD onset.

### Event-by-event image analysis of button press and SSVEP-SNR

Due to variations in the frequency and duration of PFI per participant, averaging data over participants is not straightforward. To resolve this, we performed image-based event-by-event analyses (Fujiwara et al., 2017) to investigate whether the amount of PFI reported may reflect changes in log(SNR). Within each participant, all PFI events were sorted in descending order based on the sum of buttons pressed at each time point, and over a 2.5 second time window (detailed below) per disappearance/reappearance event. For this analysis, we counted three button presses as 3 even though participants might have tried to press 4 buttons. For PFI disappearances and reappearances, we averaged this over [0, +3] seconds and [-3, 0] seconds with respect to the button press or release, respectively. We call this sum of the number of buttons pressed over these time periods “the amount of PFI”. We then resampled along the trial dimension to 100 samples to map from 0 to 1 (normalized event count) for each participant. Participant data was then smoothed along the normalized trial dimension and averaged across participants, to visualize the time-course of SNR as a function of normalized PFI. This resampling, smoothing and averaging process performed on button press responses was repeated for the event-by-event time course of log(SNR), with the order of events predetermined by the corresponding button press responses per participant. A schematic pipeline for this entire procedure is displayed in Figure 3.

**Figure 3.**
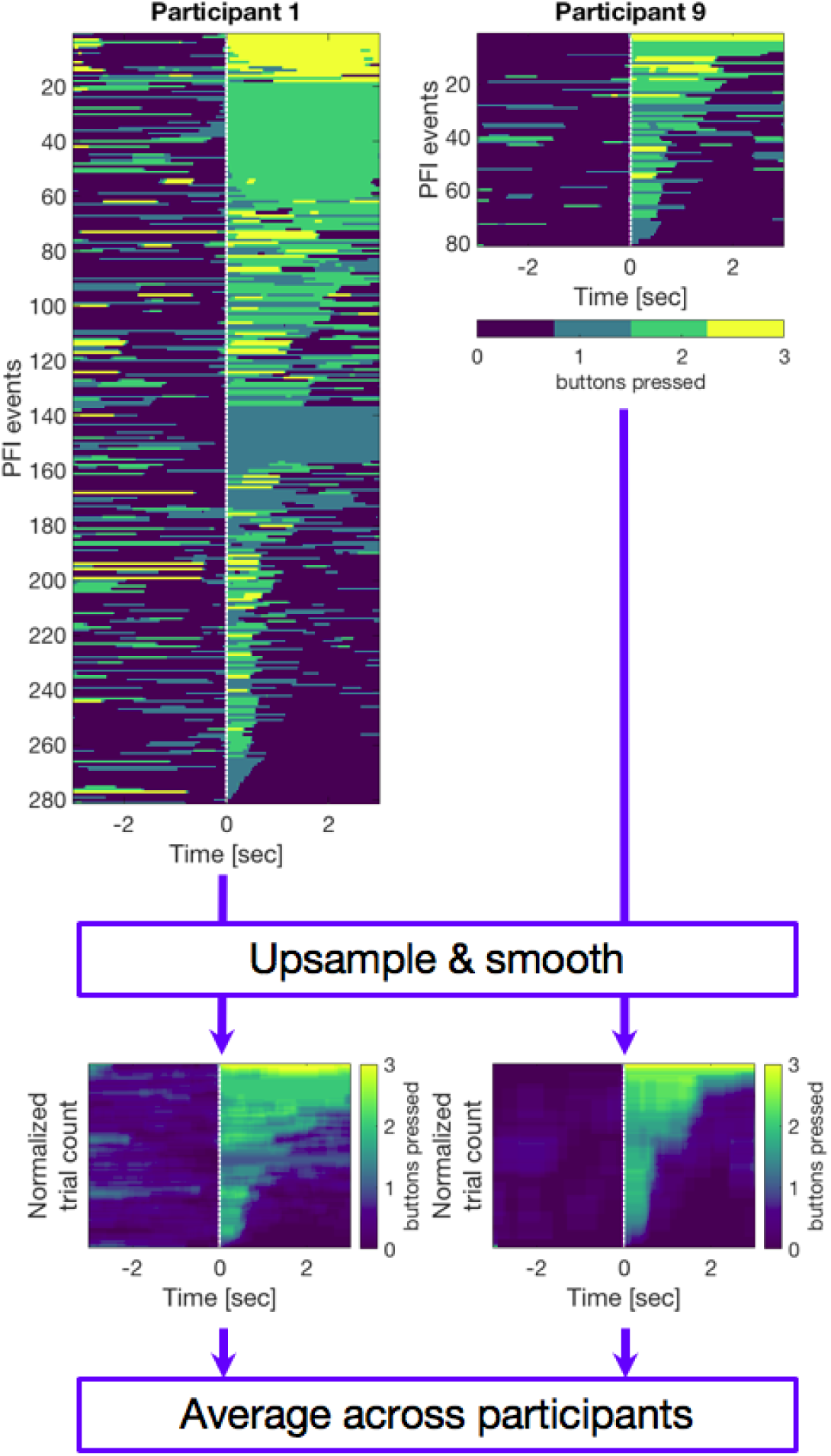
Pre-processing for event-by-event based image analyses. PFI events were first sorted according to “the amount of PFI” (the sum of buttons pressed over 2.5 seconds) occurring after a button press, or before a button-release event. Each image along the y-axis was then resampled to normalize the trial number into arbitrary units of 100 samples. A 15-sample moving average was then applied to smooth each image along the normalized event-dimension, before averaging across participants. The same process was also applied to RESS log(SNR) after sorting by the amount of PFI per event based on button press (or release) events. This image-based analysis enables us to compare PFI dynamics despite differences in the number of PFI events per participant.

To quantify the relationship between log(SNR) and the amount of PFI, we grouped events when the amount of PFI was between 0 and 1, 1 and 2, or greater than 2. A median split based on the amount of PFI resulted in similar data and subsequent conclusions. The results of this event-by-event image analysis are displayed in Figure 8.

### Reconstruction analysis to compare the impact of multiple-target disappearances and reappearances on SNR, during PFI and PMD periods

Due to our novel task design which employs multiple targets, it is necessary to account for whether the temporal dynamics of log(SNR) during PFI or PMD periods differ due to the involvement of unique mechanisms, or are attributable to differences in the requirements to report on overlapping events. For example, in our data the increase of log(SNR) at PFI onset appears to be transient, while decreases upon PFI offset are more sustained (Figure 8). Could these differences be due to the overlapping influence of temporally proximal PFI events? This is particularly important as the temporal profile of PFI and PMD may differ based on the way we programmed PMD periods; PFI events can accumulate for multiple targets in close temporal proximity, yet multiple PMD periods share a common temporal onset, even when spatially distributed. We approached this problem by performing a SNR reconstruction procedure, to model and compare the expected temporal dynamics of log(SNR) during PFI and PMD, when accounting for accumulating disappearance and reappearance events. This analysis progressed through three steps (Figure 4).

**Figure 4.**
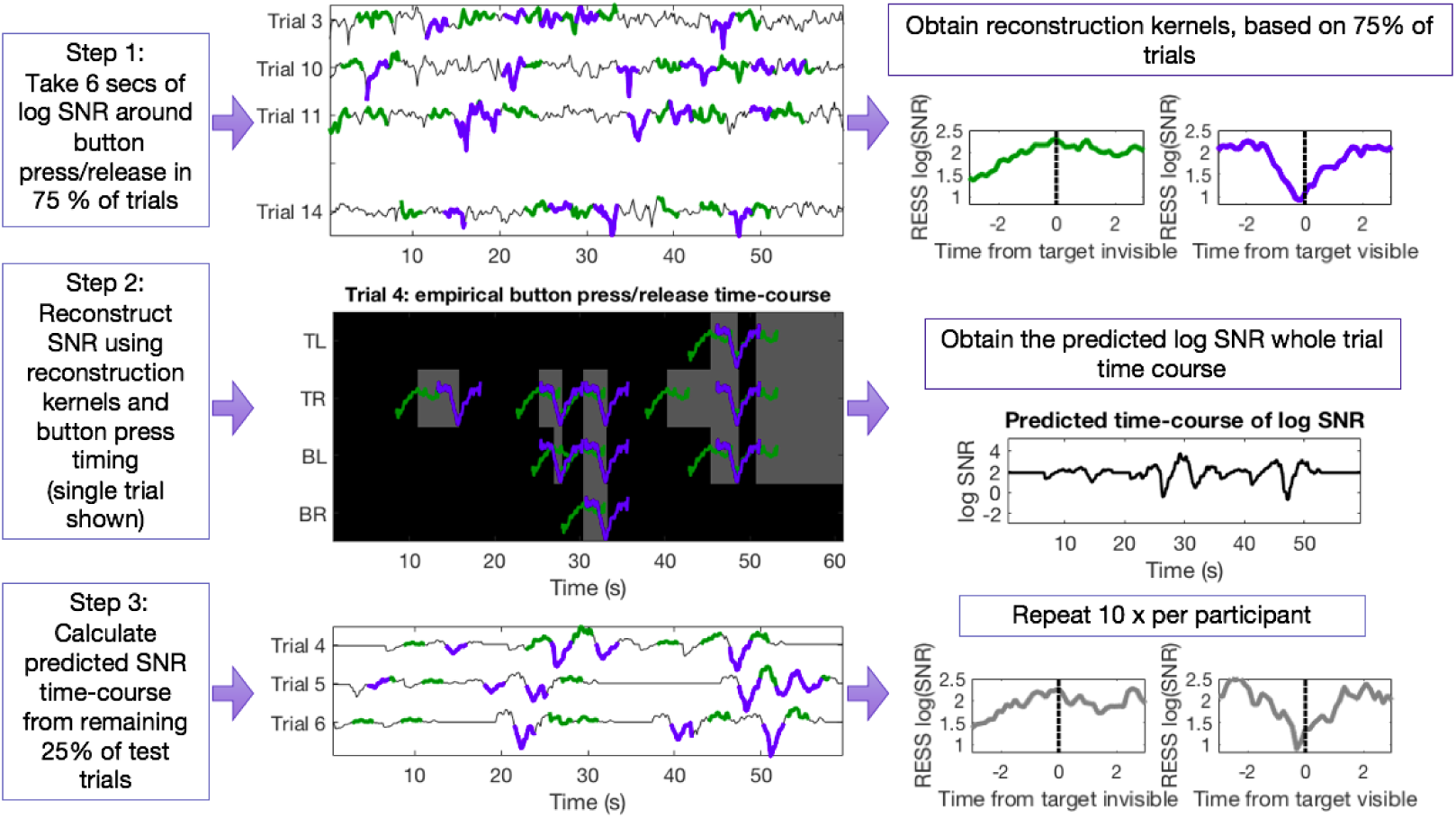
Pipeline for SNR reconstruction analysis to estimate the impact of accumulated PFI disappearances/reappearances on the observed time course of log(SNR). Step 1: we first calculated the reconstruction kernels in response to target disappearance and reappearance events from 75% of training trials per participant. Log(SNR) around button press/release events (epoched −3 to +3 seconds) is shown in green/blue, respectively. Reconstruction kernels are computed as the mean log(SNR) time course around button press/release events (over 18 trials, for this participant who had no rejected trials). Step 2: to predict the time course of log(SNR), we convolved the reconstruction kernels from Step 1 with recorded time of button press and release events in the remaining test trials (here only displaying 1 trial for demonstration purposes). As multiple PFI disappearances and reappearances can happen across target locations in close temporal proximity (< 1 second), this analysis enabled an estimation of the impact that consecutive PFI events have on SNR time course. The predicted time courses (grey) are computed as the mean log(SNR) during PFI events for test trials (over 6 trials for this participant). The predicted time courses are compared with the observed time courses from the same test trials (6 trials). This entire procedure was repeated 10 times per participant to obtain the mean predicted and observed time course for correlation analysis.

First, we calculated the mean log(SNR) time-course for PFI disappearances and reappearances using 75% of trials. Within these trials, we stepped through each time-point in the accumulative button press responses (0-3 buttons pressed), and epoched the log(SNR) time-course from −3 to +3 seconds around the time of PFI events, which we defined as any change in button press state (6-second epoch). For this analysis, we did not distinguish the number of disappearing targets at each time-point, just the direction of change (disappearing or reappearing), and obtained the mean disappearance/reappearance time courses which we subsequently used as reconstruction kernels. Second, using these reconstruction kernels, we then predicted the SNR in the remaining 25% of 60-second test trials. We did this by assuming linearity and time invariance in PFI responses, and predicted the 60-second whole-trial SNR time course by convolving the 6-second reconstruction kernels with the actual button press or release event times in the test trials. Outside of button press periods, we set the default SNR value as the baseline SNR value from the same trial (e.g. RESS log(SNR) = 2.1 above). Third, from the reconstructed 60-second time course of SNR, we epoched from −3 to +3 seconds around the PFI events and obtained the mean predicted log(SNR) time course. Figure 4b shows this procedure for one 60-second trial. We reconstructed a mean predicted SNR from across test trials, separately for PFI disappearance and reappearance. We repeated this reconstruction 10 times to obtain the mean predicted SNR per participant, which we then averaged across participants. We compared this measure to the observed mean log(SNR) time course from the same test trials.

We repeated the same procedure to compare the predicted SNR from PFI reconstruction kernels to the observed SNR during PMD periods. This was necessary due to the embedding of PMD periods within multi-target PFI, as PMD periods would often overlap with ongoing button press and -release events signifying genuine PFI. We were then able to statistically determine whether the SNR time courses during subjective and physical target disappearances/reappearances were statistically distinct, by convolving the reconstruction kernels based on (training) genuine PFI with the button press or -release event times of (test) genuine PFI and PMD periods.

To compare the predicted and the observed SNR time course, we evaluated the degree of correlation between them over the 6 seconds surrounding button press and release, obtaining R^2^ for each individual participant. For the statistical analysis, we used repeated measures two-way ANOVA, testing the main effects of background harmonics (1f = 20Hz vs 2f = 40Hz) and the nature of disappearance/reappearance (PFI vs PMD) on the R^2^ between the observed and the predicted SNR time course. The results of this reconstruction analysis are displayed in Figure 9, and comparison of model fit is shown in Tables 1 and 2.

**Table 1.**
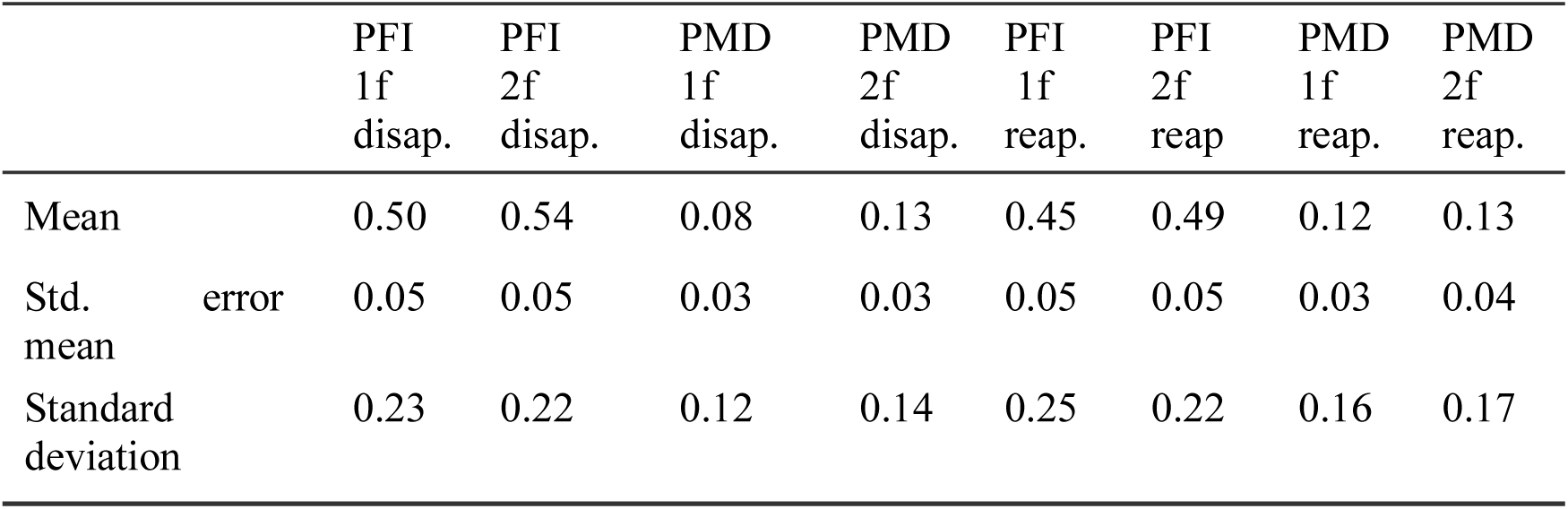
Prediction accuracy (as R^2^) across reconstruction analyses

**Table 2.** Results of 2 x 2 x 2 repeated measures ANOVA on R^2^ values

| | Sum of Squares | df | Mean Square | F | p | partial $\eta^2$ |
| --- | --- | --- | --- | --- | --- | --- |
| PFI vs. PMD | 6.24 | 1 | 6.24 | 151.01 | <.001 | 0.88 |
| Residual | 0.87 | 21 | 0.04 |  |  |  |
| 1f vs. 2f | 0.06 | 1 | 0.06 | 0.94 | 0.342 | 0.04 |
| Residual | 1.25 | 21 | 0.06 |  |  |  |
| Disap. vs. Reapp. | 0.01 | 1 | 0.01 | 0.45 | 0.512 | 0.02 |
| Residual | 0.67 | 21 | 0.03 |  |  |  |
| (PFI vs. PMD) x (1f vs. 2f) | 0.00 | 1 | 0.00 | 0.07 | 0.792 | 0.00 |
| Residual | 0.65 | 21 | 0.03 |  |  |  |
| (PFI vs. PMD) x (Disap. vs. Reapp.) | 0.05 | 1 | 0.05 | 5.34 | 0.031 | 0.20 |
| Residual | 0.21 | 21 | 0.01 |  |  |  |
| (1f vs. 2f) x (Disap. vs. Reapp.) | 0.01 | 1 | 0.01 | 0.47 | 0.499 | 0.02 |
| Residual | 0.39 | 21 | 0.02 |  |  |  |
| (PFI vs. PMD) x (1f vs. 2f) x (Disap. vs. Reapp.) | 0.00 | 1 | 0.00 | 0.28 | 0.603 | 0.01 |
| Residual | 0.34 | 21 | 0.02 |  |  |  |

Note. Type 3 Sums of Squares

### Analysis of SNR timing differences

Past research on binocular rivalry has indicated that perceptual alternations between frequency-tagged stimuli are captured in the time course of SNR, and that the time point shortly after when two SNR time courses crossover concurs with button presses to indicate a change in perception (Brown & Norcia, 1997; Jamison, Roy, He, Engel, & He, 2015; Tononi & Edelman, 1998; Zhang et al., 2011). We were interested to see whether changes in SNR could also predict button presses/releases in our multi-target PFI paradigm, and compared the onset at which 1f and 2f responses indicated a change in perception. For background related SNR, this requires overlaying the same SSVEP response (e.g., 20 Hz) during both disappearance and reappearance, as opposed to comparing two unique simultaneous SSVEPs as is usually done in binocular rivalry research (e.g. Zhang et al., 2011; Katyal et al., 2016).

Initial data inspection revealed that at the participant level, these overlaid SNR time-courses could cross multiple times during PFI events. As a result, rather than using an arbitrary SNR cross-over point as the focus of our analysis, we compared the disappearance and reappearance time courses using paired samples *t*-tests at each time point. Clusters of significant time points were identified which satisfied *p* < .05 (uncorrected) over a minimum of 300 ms, a time window which corresponds to two adjacent time points in our moving-window SNR. Per participant, the first time point in these clusters, which occurred after the time point where the two time courses crossed each other, was taken as the earliest time point at which the SNR differentiates between target disappearance and reappearance in our analysis. We then compared the earliest time-points detected by this procedure for 1f and 2f, using a two-tailed paired-samples t-test. We also performed the same analysis to compare the time course of SNR during physical target disappearance and reappearance due to PMD periods. Note that this cluster-procedure is distinct from the procedure described below, which retains the largest cluster for non-parametric tests (*Statistical analysis - EEG*).

### Spatial correlation analysis

To perform the spatial correlation analysis, we calculated the time-course of a 64-channel correlation between 1f and 2f log(SNR). Due to differences in the number of PFI events and PMD periods, we down-sampled (with replacement) the number of PFI events to 24, which was the maximum number of available PMD periods. We then calculated the correlation for this subset of trials, and repeated this analysis 100 times to obtain a distribution of down-sampled correlation values. The mean correlation value from this down-sampled distribution was then used to compare the spatial correlation of PFI and PMD periods. The results of this analysis are displayed in Figure 11.

### Statistical analysis – EEG

To assess the significance of SSVEP peaks in the EEG spectra, we corrected for multiple comparisons with a False Discovery Rate (FDR) of .05 (Benjamini, Krieger, & Yekutieli, 2006; Yekutieli & Benjamini, 2001). For corrections of multiple comparisons on the time courses, we used temporal cluster-based corrections (Davidson, Alais, van Boxtel, & Tsuchiya, 2018; Maris & Oostenveld, 2007). For this analysis, the sum of observed test-statistics (e.g., *t* scores) in a temporally contiguous cluster were retained for comparison with a permutation-based null distribution. Specifically, first, we detected any temporally contiguous cluster by defining a significant time point as *p* < .05 uncorrected (Maris & Oostenveld, 2007). Then, we concatenated the contiguous temporal time points with *p* < .05 and obtained a summed cluster-level test statistic for the cluster. Second, we repeated this procedure after shuffling the subject specific averages within each participant 2000 times. From each of the 2000 shuffled data, we obtained the summed cluster-level test statistics at contiguous temporal time points with *p* < .05 uncorrected, which served as a null distribution. We regarded the original observed effect to be significant if the original summed cluster-level statistics exceeded the top 97.5% of the null distribution of the summed statistics (as *p_cluster_* < .025).

## Results

### Overview

Our presentation of the results will be structured as follows. First, we confirmed that our overall SSVEP frequency-tagging was successful (Figure 5). Second, we checked if the behavioural reports during PMD periods were correlated with neural activity (RESS log(SNR), Figure 6). Third, we investigated the behavioural reports during genuine PFI events, and focused on whether or not spatially separated PFI targets interact across visual quadrants (Figure 7). Fourth, we then focused on RESS log(SNR) during PFI events, testing if the amount of PFI correlated with the strength of frequency-tagged EEG activity induced by our flickering background (Figure 8). Fifth, we devised an SNR reconstruction analysis to estimate the influence of multiple PFI events in close temporal proximity on the RESS log(SNR) (Figure 9). Sixth and finally, we also found unexpected temporal (Figure 10) and spatial (Figure 11) differences between PFI events and PMD periods, with respect to the first (1f) and second harmonic (2f) responses to background flicker, which we interpret in our Discussion.

**Figure 5.**
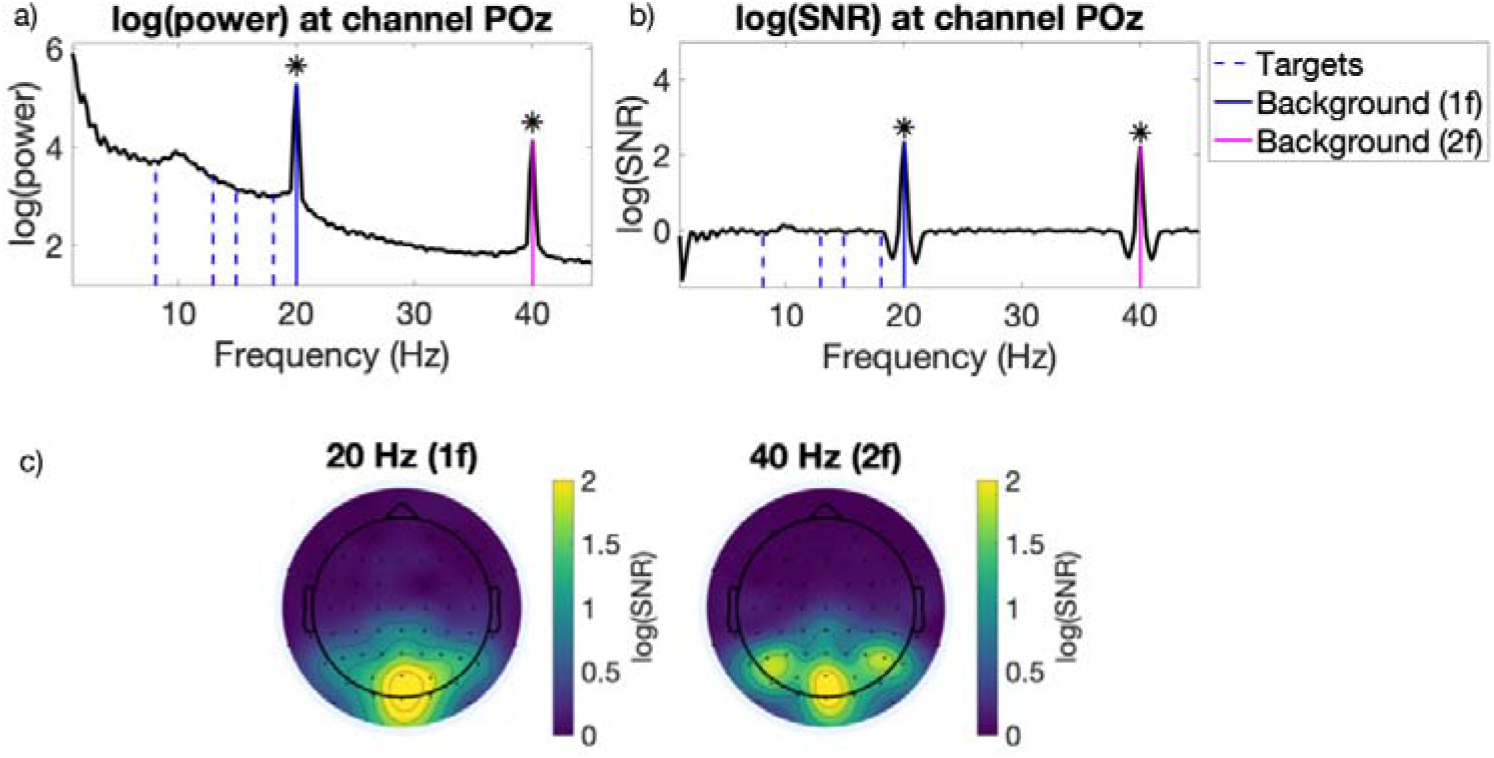
Average SSVEP responses in our paradigm. a) The mean log(power) spectrum, and b) log(SNR) over all participants and periods of PFI at channel POz. Asterisks mark peaks significantly different from 0, FDR-adjusted across all frequencies to *p* < .05. c) Topoplots for the mean log(SNR) at 20 Hz (1f; stimulus flicker), and 40 Hz (2f; stimulus harmonic) for background-related SSVEPs. The mean is taken across participants over all epochs, excluding PMD periods.

**Figure 6.**
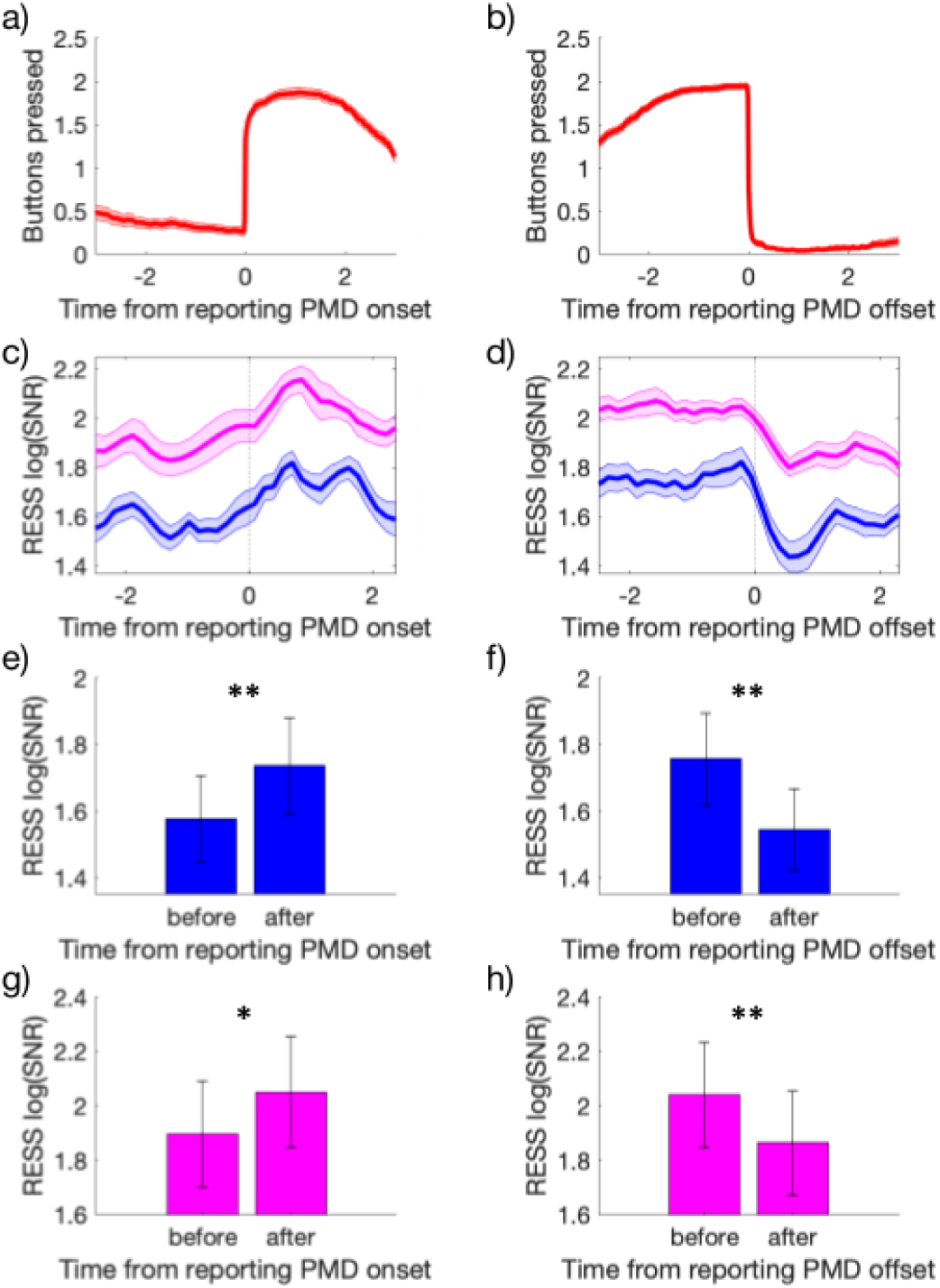
Button press time course and background RESS log(SNR) around PMD periods. a-b) mean (± 1 SEM) button press time course across participants when responding to the physical removal of targets near the onset (a) and the offset (b) of PMD periods. c-d) RESS log(SNR) for background SSVEP at 1f (20 Hz; blue) and 2f (40 Hz; magenta). Shading represents ± 1 SEM corrected for within participant comparisons Cousineau, 2005). e-h) Bar-charts for the statistical comparisons reported in text, comparing RESS log(SNR) before and after button press during PMD periods.

### Successful frequency-tagging of dynamic background in PFI display

We first investigated the log(SNR) of target and background flicker frequencies and their harmonics. Using a short window (2.9 second duration, see Methods), we found strong and occipitally localized responses to background flicker, but no clear responses to target flicker. Even when we analysed the data with the longest time window (60 second for one trial, including PMD periods) with the highest frequency resolution, we did not detect reliable target-related SSVEPs (Figure 5a), owing to their small size and eccentricity.

While the 1f (20 Hz) and 2f (40 Hz) frequency-tagged responses to our background display were strongest at POz, the spatial topographies differed between 1f and 2f (Figure 5b). The 1f response was localized to midline occipital electrodes, while the 2f response extended beyond these regions to include lateral parieto-occipital and parietal electrodes. We continued to analyse the time-course of log(SNR) for background-related 1f and 2f responses after applying rhythmic entrainment source separation (RESS; Cohen & Gulbinaite, 2017), to optimally extract the SNR per participant given these differences in source topography and to avoid multiple comparisons across electrodes (see Methods). From here, all SNR values we present are the RESS log(SNR) (except for the spatial correlations presented in Figure 11).

### Frequency-tagging during PMD periods

Having identified the successful entrainment of background responses (Figure 5), we analysed the time course of changes to the RESS log(SNR), focusing on 1f and 2f during PMD periods. As SSVEPs tend to be weak for peripherally presented stimuli Norcia et al., 2015), we checked if the physical removal of targets was strong enough to alter the time course of the SNR activity. During PMD periods, we compared the mean RESS log(SNR) during −2 to −0.1 to +0.1 to +2 seconds (two-tailed paired samples *t*-tests). The SNR to background flicker increased upon target removal (1f, *t*(21) = 3.80, *p* = .0011; 2f, *t*(21)= 2.21, *p* = .038). The background SNR also decreased upon target return (1f, *t*(21)= −3.51, *p* = .0021; 2f, *t*(21) = −3.50, *p* = .0021). The increase/decrease of the RESS log(SNR) started upon button press/release, which we return to and investigate in our SNR-reconstruction analysis (Figures 4 and 9). These results are consistent with an interpretation that the background 1f and 2f SNR increases when peripheral regions are physically interpolated by the flickering background display.

### Synergistic effect of multi-target PFI

Next, we turn to the behavioural analysis of the genuine PFI events before interpreting the EEG effects. Specifically, we investigated whether our unique multi-target design had captured an interaction between the four simultaneously presented peripheral targets. Previous research has suggested that neighbouring targets within a single visual quadrant may disappear together (De Weerd et al., 1998). Our design allowed us to examine whether much more widely distributed peripheral targets also interact. Such an interaction would be non-trivial if occurring across the disparate retinotopic locations of all four quadrants of the visual periphery, and would imply the involvement of potentially high-level neural mechanisms that have access to these long-range relations (Wagemans et al., 2012).

First, we analysed whether the number of targets simultaneously invisible were related to 1) the number of PFI events per trial, 2) the average duration of PFI per event, and 3) the total duration of PFI per trial (Figure 7, blue bars). In theory these three variables can vary independently, and in practice they can dissociate (Bonneh et al., 2014; McEwen et al., 2018; Thomas et al., 2017). While periods when all targets were visible had the longest average duration and total duration (i.e., the number of invisible targets = 0), the more interesting trends were found as the number of invisible targets increased. While simultaneous disappearances of 3 or 4 targets were rare (only 2.9 events per trial; Figure 7a), when they happened, the event tended to be sustained for a long duration (2.2 sec, Figure 7b). As a result, the total duration of 3 or 4 target invisibility (7.6 seconds per trial, Figure 7c) is comparable to that of 2 target invisibility and longer than that of 1 target invisibility, which happened at the highest rate (8.8 events per trial, 5.8 seconds in total per trial). We formally tested this linear trend by LME analysis and likelihood ratio tests (see Methods). The number of invisible targets (nPFI) significantly affected 1) the number of PFI events per trial (χ^2^(2) = 47.83, *p* = 4.1×10^-11^, 2) the average duration of PFI per event (χ^2^(2) = 23.59, *p* = 7.53×10^-6^) and 3) total PFI duration per trial (χ^2^(2) = 7.27, *p* = .026).

**Figure 7.**
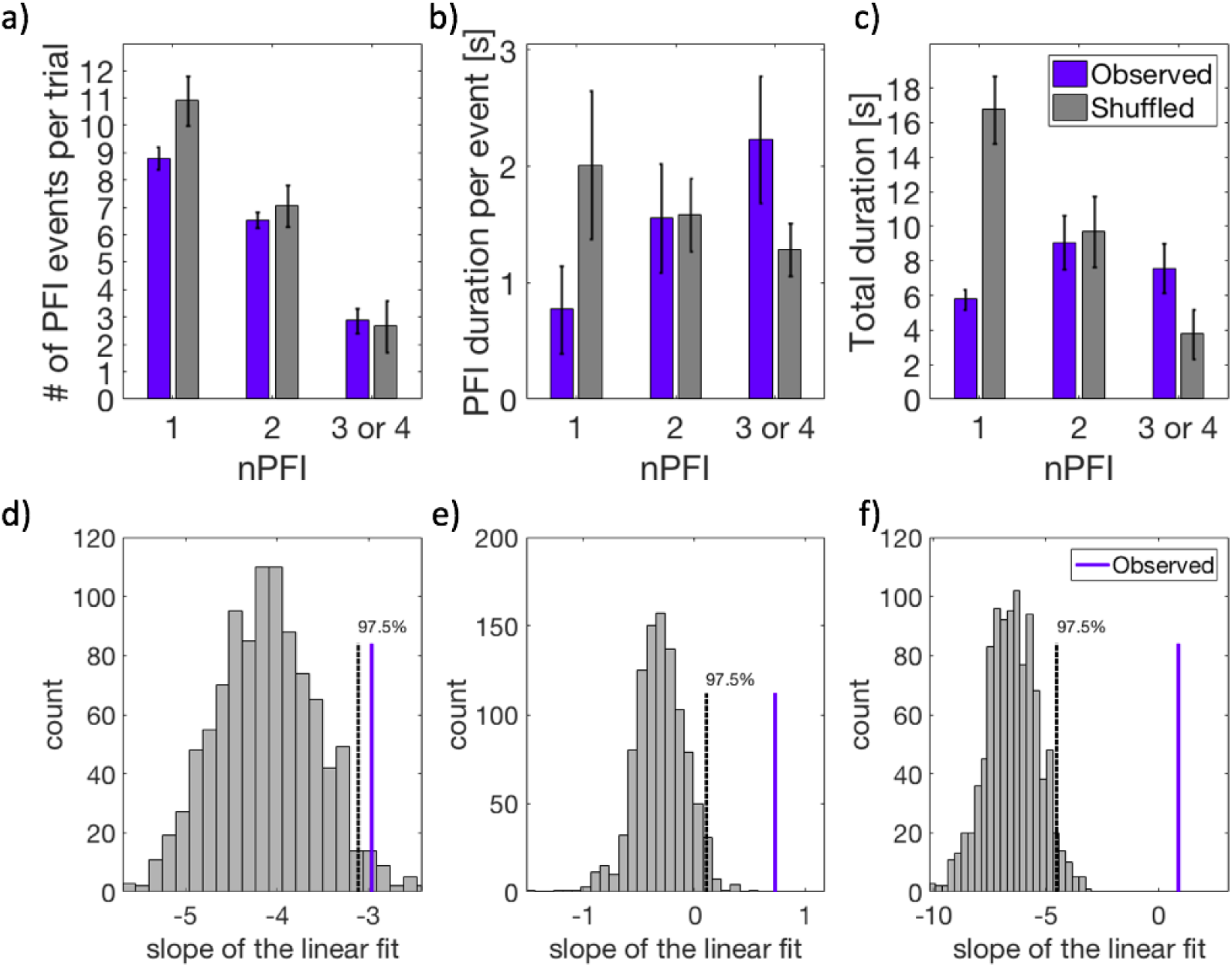
Behavioural data. a) The number of PFI events per trial, b) the mean duration per PFI event, and c) total duration of PFI per trial, as a function of the number of invisible targets (nPFI). All panels display both observed (blue) and shuffled (grey) data. For the observed data, error bars represent 1 SEM, corrected for within-participant comparisons(Cousineau, 2005). For the shuffled data, we first computed the SEM within each shuffled data set across participants. Then, as the error bar, we show the mean of the SEM across 1000 shuffled sets. d-f) Slope of the linear fit for each of the PFI variables in a-c as a function of nPFI for the observed (blue line) vs the shuffled data (1000 sets, grey histogram).

These significant trends imply that interactions among distant targets occur in a synergistic way, and that when one target is invisible it is often accompanied by other invisible targets. To directly test if this is the case, or if these trends occur by chance, we employed a shuffling analysis (see Methods). For this, we first sub-selected the button press time course for each location from any four trials (with replacement) and re-computed the behavioural analysis per participant. We repeated this shuffling procedure 1000 times, and from each shuffled dataset we retained the mean PFI data across participants. As the location of each button press in shuffled data could come from any independent trial (e.g., top left = trial 1, top right = trial 23, bottom left = trial 18, bottom right = trial 18), this shuffling procedure conserved the mean number of PFI events overall, while estimating the level of simultaneous invisibility between multiple PFI targets that occurs by chance, when locations are independent.

In the shuffled data, the number of PFI events per trial decreased as the number of invisible targets (nPFI) increased, which is similar to what we observed in the empirical data (10.8, 7.1, and 2.7 events per trial for 1, 2, and 3 or 4 target invisibility; Figure 7a, grey bars). However, the trend for shuffled data was quite different from the empirical data for the average durations per PFI event, which were roughly equal across nPFI in shuffled data (2, 1.6, and 1.3 seconds, respectively; Figure 7b), and the total duration of PFI per trial, which decreased as a function of the number of invisible targets (16.6, 9.7, and 3.8 seconds, respectively; Figure 7c) .

To statistically evaluate these trends between the observed and the shuffled data, we compared the slopes of the linear fit (LME, with random intercepts for each subject) for each of the three PFI variables as a function of the number of invisible targets (nPFI; 1, 2, 3 or 4). For all variables, the observed slope was outside the top 97.5% of the slopes in the shuffled data (corresponding to two-tailed *p* < .05, Figure 7d-f). Notably, Figure 7e and f establish that the observed positive slope for observed data in Figure 7b and 7c are contrary to the expected negative slope in shuffled data. In other words, if there are no spatial interactions between distant targets, as in our shuffled data, then we should expect the simultaneous invisibility of 3 or 4 targets to be highly unlikely, and sustained for a shorter duration. By contrast, the observed data show that as more targets are involved with a disappearance event, the longer the disappearances are sustained, strongly suggesting a facilitative interaction between invisible peripheral targets. We return to this synergistic effect of multi-target PFI in our Discussion.

### SSVEP time course: event-by-event image analysis reveals graded changes in conscious perception

After demonstrating that spatially distributed targets were interacting, strongly implying the involvement of high-level neural mechanisms during PFI, we turned to the neural correlates of PFI via EEG analysis of SSVEPs. We first visualized how the changes in PFI were related to changes in the log(SNR) of background flicker using an event-by-event image-based analysis. To compare the time course of button press and SNR across participants, we first sorted, per participant, all instances of PFI disappearance (or reappearance) by the sum of the number of buttons simultaneously pressed over 2.5 seconds after (or before) the button press, which we define as “the amount of PFI” (see Methods and Figure 8). We then resampled each participants image into a uniform height, to obtain the across-participant mean despite the differences in individual PFI dynamics (see Methods and Figure 3). This results in the highest (and lowest) rows of the figures representing events with the highest (and lowest) amount of PFI (Figure 8a-b). Figure 8c-f show the corresponding RESS log(SNR) related to 1f and 2f background SSVEP responses.

**Figure 8.**
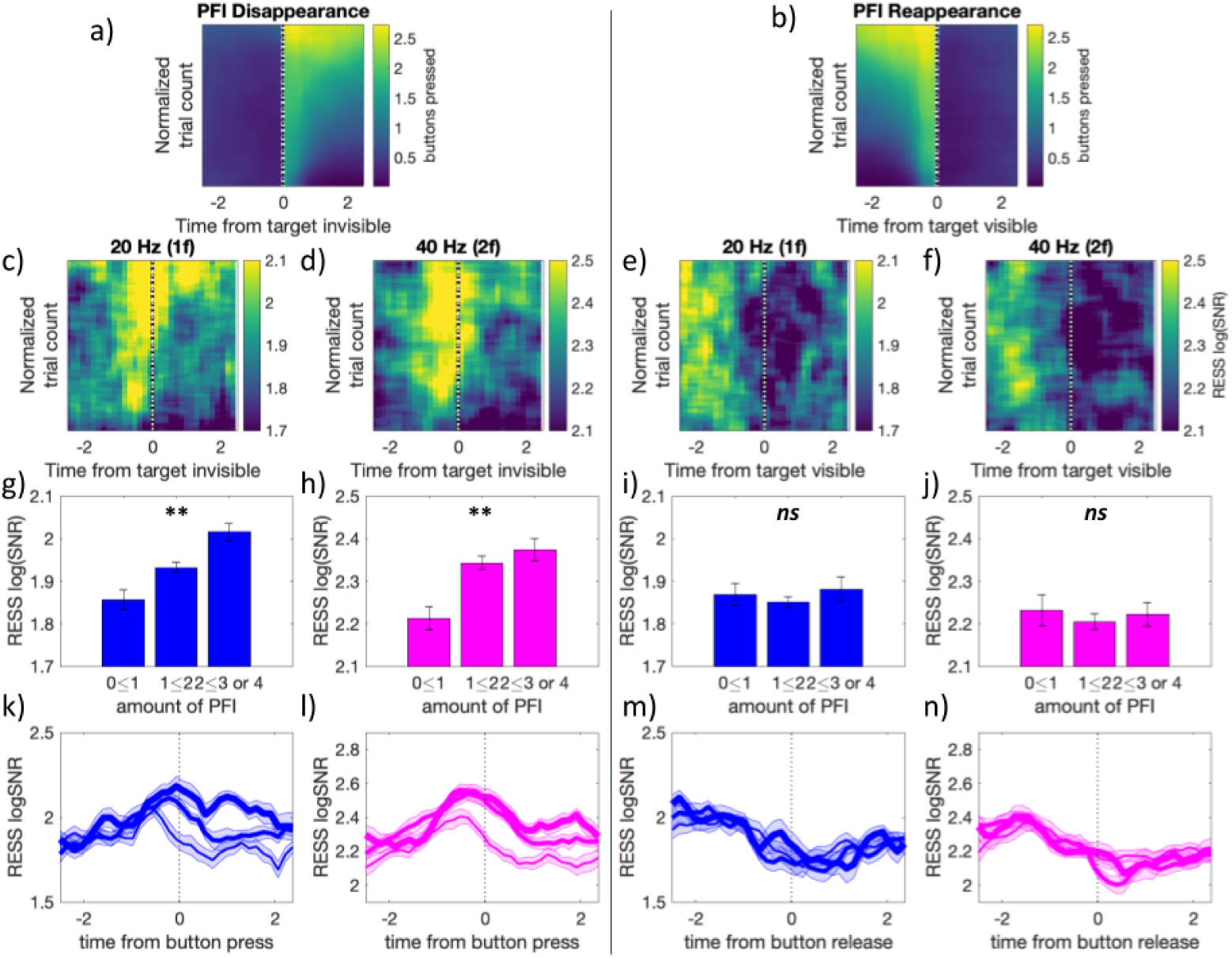
The amount of PFI is correlated with the RESS log(SNR) around PFI disappearances, but not reappearances. Event-by-event image analysis of button press and release (a and b) and RESS log(SNR) (c-f) after sorting based on the amount of PFI per event, per participant. Background responses at 1f are shown in blue and those at 2f are in magenta. g-j) Bar graphs for the mean RESS log(SNR) over −2.5 to 2.5 seconds as a function of the amount of PFI. k-n) The time course of RESS log(SNR) around the button press or release, separated by the amount of PFI, with three levels of the amount indicated by the thin, middle and thick lines. Error bars in g-j and shading for k-n indicate 1 SEM across participants (adjusted for within-participant subject comparisons (Cousineau, 2005).

From this analysis, two qualitative insights emerged. First, that RESS log(SNR) for 1f and 2f increase just before button press when targets disappear (at time = 0), and increase with the amount of PFI (Figure 8c-d). Second, RESS log(SNR) for 1f and 2f decrease just before button release at target reappearance, but there is no dependence on the amount of PFI (Figure 8e-f).

To quantitatively compare these differences, we split SNR time courses based on the amount of PFI. Figure 8g-j show the mean RESS log(SNR) over each 6 second epoch, separately averaged for events with the amount of PFI between 0 and 1, 1 and 2, or greater than 2. Around the target disappearance events, we found a significant linear effect for the amount of PFI on the SNR for both 1f (χ^2^(1) = 8.75, *p* = .003) and 2f (χ^2^(1) = 8.21, *p* = .004) responses to background flicker (Figure 8g-h). Around target reappearance events, by contrast, the amount of PFI did not significantly affect the SNR (Figure 8i; *p* = .76; Figure 8j; 2f; *p* = .83). Figure 8k-n displays the time course of the SNR separately for each level of the amount of PFI around the time of button press and release.

### Reconstruction analysis: SNR time courses during PFI are distinct from those in PMD periods

The previous analysis has shown that changes to the log(SNR) of background flicker were related to the amount of PFI, we next considered whether these changes could be distinguished from the changes evoked by PMD periods. Contrasting PFI and PMD could be used to isolate the neural substrates which are unique to an endogenous change in perception, as opposed to a physically induced change.

However, to do this, we first need to take into account the effects of closely spaced or overlapping button responses that are required in our multi-target PFI task. Unlike other tasks that have investigated the neural correlates of multistable perception with a single target, our task design allowed graded changes in consciousness to occur in close temporal proximity (< 1 second), and even to overlap (Figure 1b). This was not the case for our programmed PMD periods, which occurred with a simultaneous onset and offset. To account for whether the changes in log(SNR) during PFI are distinct from those during PMD, we performed an SNR-reconstruction analysis. This analysis quantified the stability of log(SNR) dynamics during PFI, and whether differences in log(SNR) during PMD could be predicted from the same data. In brief, we used 75% of trials to construct reconstruction kernels, which were the changes to log(SNR) during PFI in this ‘training’ data. We then applied these kernels to the remaining 25% of ‘test’ trials, aligning each kernel to the recorded button press time-points (see Methods and Figure 4). We then compared the predicted time course of log(SNR) based on reconstruction kernels, with the actual time course in the test trials during genuine PFI and PMD periods. Figure 9 visualizes the high quality of prediction for the genuine PFI (Figure 9e and g) and the poor predictive quality for PMD periods (Figure 9f and h). These differences in predictive accuracy of our SNR reconstruction analysis implicate distinct neural substrates for PMD and PFI.

**Figure 9.**
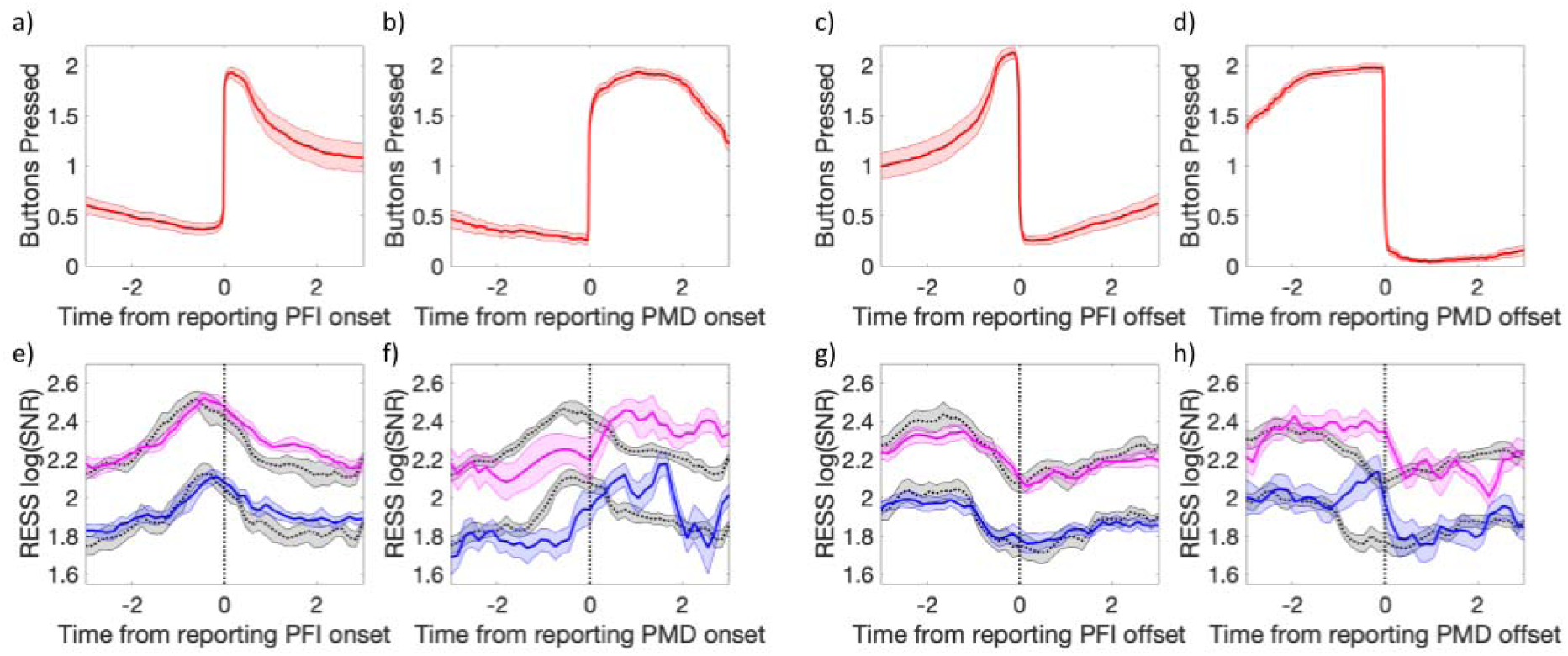
Reconstruction analysis. a-d) mean button press and e-h) RESS log(SNR) time course across participants around genuine PFI events for disappearance (a, e) and reappearance (c, g) and around PMD disappearance (b, f) and reappearance (d, h). Note that for all panels, time 0 is always defined by a button press or release. e-h) The observed SNR time course is shown from test trials (blue for 1f and magenta for 2f), which were not used to construct the reconstruction kernels. The correlation (R^2^) between the observed SNR and the predicted SNR (shown in grey) was used to quantify prediction accuracy. Shading represents 1 SEM across participants (corrected for within participant comparisons; (Cousineau, 2005).

We quantified reconstruction prediction accuracy as the degree of correlation between the predicted and the observed time course. We calculated R^2^ between the respective 6-second RESS log(SNR) around button press/release events during genuine PFI and PMD periods. For both 1f and 2f, the predicted SNR was correlated more strongly with genuine PFI than the PMD, for both disappearances and reappearances (Table 1). Using 3-way repeated measures ANOVA (Table 2), we confirmed that the prediction accuracy is significantly better for the genuine PFI than PMD periods (main effect: *F*(1, 21) = 151.01, *p* = 4.7 x 10^-12^). We found no or weak main effects or interactions due to other factors (i.e., 1f vs 2f, disappearances vs reappearances).

### Timing differences: 1f and 2f background-related responses are temporally distinct during PFI

Our reconstruction analysis similarly predicted both the 1f and 2f components of background-related SNR during PFI events, which is not surprising given that these responses were driven by the same stimuli. Curiously, however, these harmonic responses were topographically distinct (Figure 5b). As there is a nascent literature suggesting that SSVEP harmonics may correspond to separate cognitive processes (Kim et al., 2007, 2011)we next investigated these spatiotemporal differences in more detail.

First, we investigated whether the RESS log(SNR) time course differed depending on the nature of disappearances/reappearances: due to physical (PMD) or perceptual (PFI). We compared the time courses between target disappearance and reappearance, superimposing the time courses for disappearances/reappearances in the same plot and calculating the periods at which the RESS log(SNR) significantly differed between them. For 1f (Figure 10a and 10b, blue), the RESS log(SNR) during disappearances (solid lines) became larger than that during reappearances (dotted lines), consistent with an increase in background-SNR during the phenomenal experience of targets becoming filled-in by the background. This effect occurred from 0.67 seconds prior to subjective report (paired *t*-tests, *p_cluster_* < .001). Notably, these effects occurred 1.06 seconds later for PMD periods (Figure 10b, from 0.39 seconds, *p_cluster_* < .001). For 2f (Figure 10a-b, magenta), the RESS log(SNR) also became larger during disappearances than reappearances from .97 seconds prior to report (*p_cluster_* < .001), and again, were shifted roughly 1.36 seconds compared to the PMD-related time course (Figure 10b, from 0.39 seconds; *p_cluster_* < .001).

**Figure 10.**
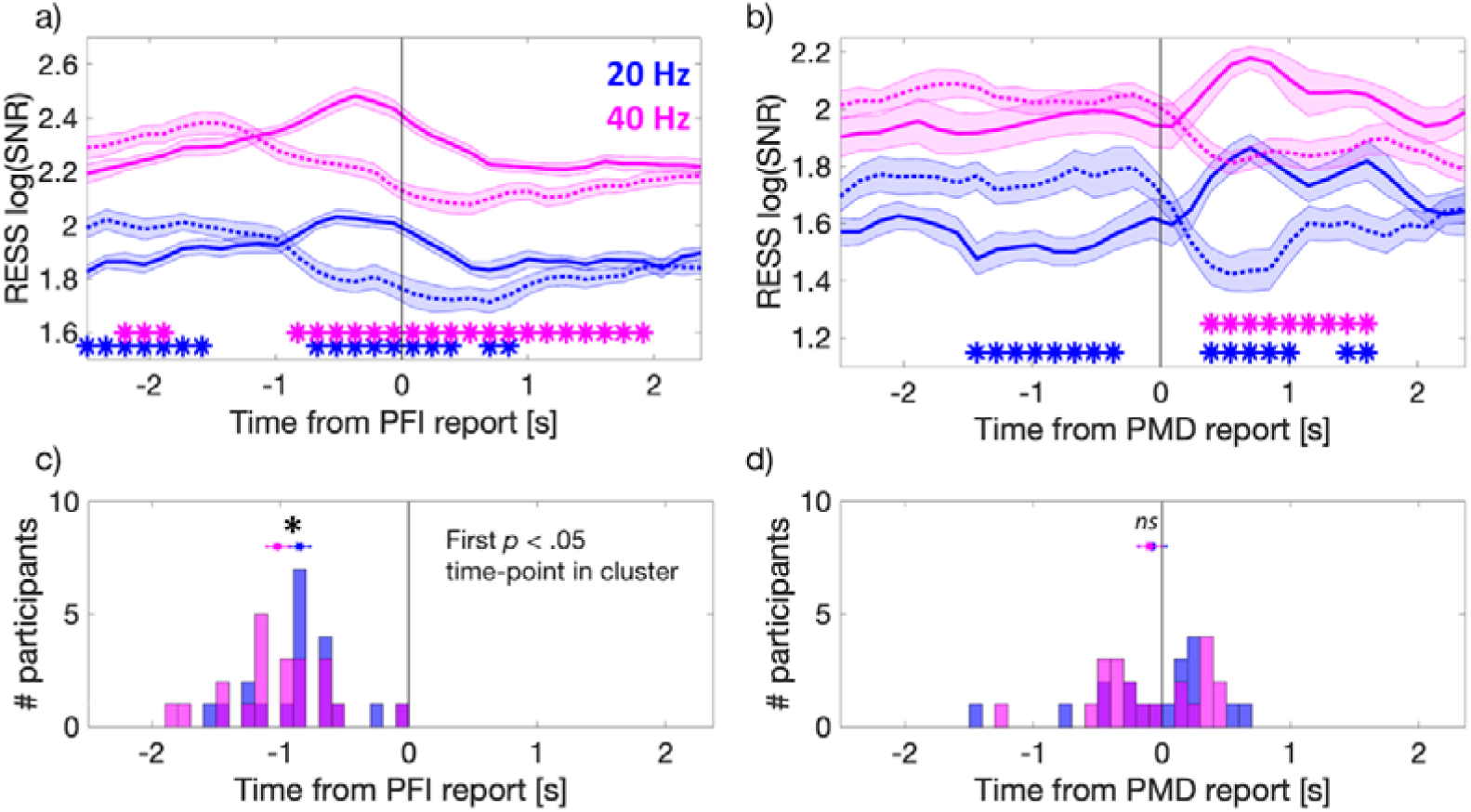
Distinct temporal profile of the harmonic responses. a and b) Group-level relative time course of the 1f (20 Hz, blue) and 2f (40 Hz, magenta) RESS log(SNR) during PFI events (a) and PMD periods (b). Solid and broken lines represent disappearance and reappearance, respectively. c and d) Participant-level histograms for the first significant time point when comparing between the RESS log(SNR) for disappearance and reappearance during PFI (c) and PMD (d). Horizontal lines indicate 1 SE about the mean corrected for within-subject comparisons (Cousineau, 2005).

The observed divergence (0.3 seconds) in the time for significant changes to 1f and 2f SNR seemed quite large given that both 1f and 2f were evoked from the same stimulus, using identical participants and events. As such we further investigated if this effect could be observed at the participant level. For this analysis, we calculated for each participant the first time point at which the strength of background RESS log(SNR) during disappearance was significantly greater than during reappearance, using running paired *t*-tests. The first time-point in a temporally contiguous cluster was retained per participant for both 1f and 2f RESS log(SNR), and then compared at the group level. Using this criterion, we found that 2f responses crossed over at −1.02 seconds (*SD* = 0.41), 170 ms seconds earlier than 1f responses, at −0.85 seconds (*SD* = 0.37, Wilcoxon signed rank test, z = 2.13, *p* = .012). No difference was observed in cross over time for the PMD-related 1f and 2f time courses (*p* = .14).

To confirm this difference was due to the latency of crossover points for 1f and 2f responses, we also performed a non-parametric jack-knife resampling procedure to estimate the standard error of the difference between crossover timepoints across subjects (Kohler et al., 2018). Specifically, we calculated the difference in 1f and 2f crossover timepoints after leaving each participant out on a single permutation. We then estimated the standard error of all crossover differences in our *N*-1 subsets over all participants (using Equation (2); of Miller et al., 1998), and then derived a *t*-statistic from this estimate (Equation (3); Miller et al., 1998). This procedure confirmed that the difference between 1f (*M=* - 0.94, *SD* = 0.018 seconds) and 2f (*M*= −1.18, *SD* =0.03 seconds) crossover points was also statistically significant (*t*(21)= −1.81, *p* = .043). The group level RESS log(SNR) time series are compared in Figure 10 a-b), with the distribution of participant-level first significant time-points displayed in 10 c-d).

### Spatial Correlation: 1f and 2f background responses are spatially distinct during PFI

One potential factor that could have contributed to the difference in the crossover time between 1f and 2f is a difference in the spatial filters used for 1f and 2f within RESS analysis. In fact, when we focused only on the (non-RESS) log(SNR) from a single electrode (POz), the difference in cross-over times between 1f and 2f was not significant at the group or participant level. Given this, we further analysed whether the spatial characteristics for 1f and 2f were also distinct without using RESS spatial filtering during PFI.

Around the PMD events, spatial correlations across 64 channels were constant (Figure 11b). However, when targets disappeared during PFI, the spatial correlation between 1f and 2f transiently increased (Figure 11a). The difference between the time courses was significant for the time-window −0.67 to 0.25 seconds around subjective report (paired *t*-tests at each time point, *p_cluster_* < .001). The same pattern of results was maintained when using a parietal or occipital sub-region of electrodes (but no change in correlation was seen for frontal or temporal electrodes), indicating that synchronous changes in predominantly parieto-occipital SNR were responsible for changes to the whole-head correlation over time. The same pattern was also observed when subtracting the mean log(SNR) per channel prior to calculating this spatial correlation over time.

**Figure 11.**
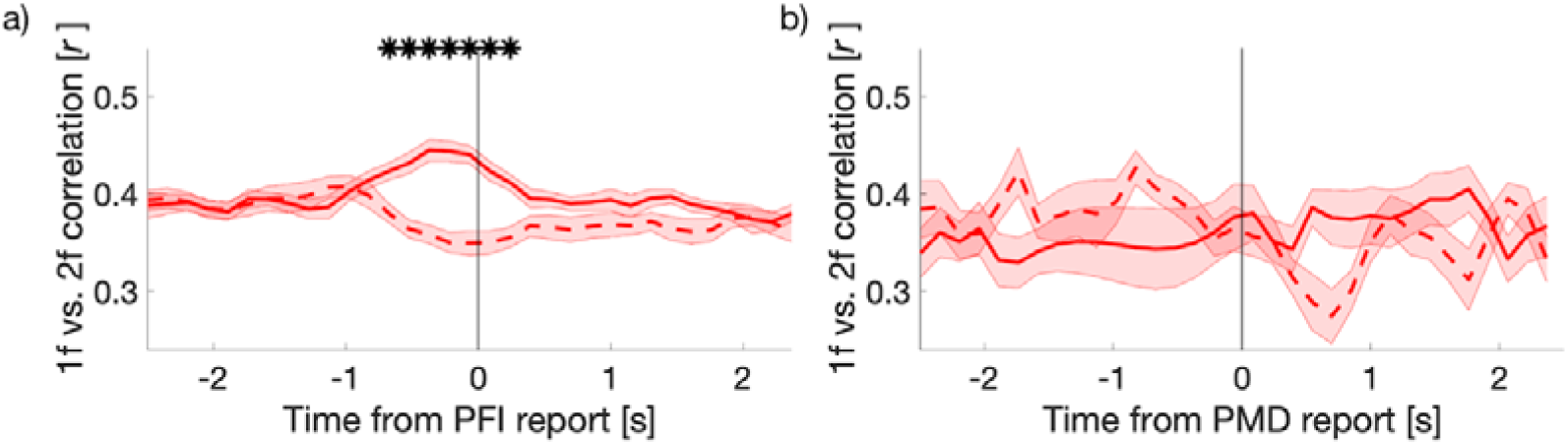
Time course of the spatial correlation coefficient (r) between 1f and 2f (non-RESS) log(SNR) across 64 electrodes. Correlation coefficient was computed across 64 electrodes at each time point per participant. The mean time courses of correlation coefficients are shown for target disappearance (solid), and reappearance (dotted) around a) PFI, and b) PMD periods. For PFI, we show the mean correlation value obtained after down-sampling PFI events to 24 (the maximum number of PMD periods), over 100 repetitions of this down-sampling procedure. Asterisks denote the time points with significantly different correlation coefficients between PFI disappearances vs reappearances (paired t-tests, cluster corrected). Shading reflects the SEM across subjects corrected for within-subject comparisons (Cousineau, 2005).

## Discussion

We combined a multi-target perceptual filling-in (PFI) paradigm with frequency-tagged EEG. This combination has revealed novel insights into the mechanisms of PFI phenomena, including unexpected asymmetric neural correlates for graded disappearances and reappearances (Figure 8), and spatiotemporal distinctions between steady-state visual evoked potential (SSVEP) harmonics (1f and 2f background responses, Figure 10 and 11). Here, we discuss these findings focusing on several advantages of our experimental paradigm.

### Multi-target PFI to track changes in conscious perception

Frequency-tagging has been used to study the neural correlates of consciousness, mainly in combination with binocular rivalry (Brown & Norcia, 1997; Jamison, Roy, He, Engel, & He, 2015; Katyal et al., 2016; Sutoyo & Srinivasan, 2009; Tononi et al., 1998; Zhang et al., 2011). When reporting on perceptual reversals in these paradigms, neural activity that is associated with purely perceptual processes has been entangled with the processes of attention and the act of report (Aru, Bachmann, Singer, & Melloni, 2012; Miller, 2007; Tsuchiya, Wilke, Frässle, & Lamme, 2015; van Boxtel & Tsuchiya, 2014; van Boxtel et al., 2010a). To reduce these confounds, replays with the physical removal or alternation of stimuli have been used as a standard control condition to compare with, for example, genuine perceptual switches in binocular rivalry (Frässle, Sommer, Jansen, Naber, & Einhäuser, 2014; Lumer, Friston, & Rees, 1998). As the requirements for both perceptual and physical reversals involve attention and report, it was hoped that contrasting these conditions would isolate the neural processes specific to endogenously generated changes in consciousness. Despite various attempts, generating physical reversals/disappearances that perceptually match endogenously-generated conscious changes in perception remains a significant challenge, due to highly complex phenomenal dynamics during rivalry (Knapen, Brascamp, Pearson, van Ee, & Blake, 2011; Wilson, Blake, & Lee, 2001). Until these report-related attentional confounds are resolved, results from such experiments, particularly binocular rivalry, need to be interpreted with caution (Blake, Brascamp, & Heeger, 2014; Frässle et al., 2014; Naber, Frässle, & Einhäuser, 2011).

Unlike binocular rivalry, perceptual changes during PFI are crisp and simple, suggesting PFI can prove to be a useful psychophysical tool to study the NCC. The simplicity of PFI phenomenology allowed us to 1) generate PMD events that were difficult to distinguish from real PFI (see Movie 1), and 2) to ask untrained participants to accurately and simultaneously report on multiple targets, while allowing us to check the quality of their report. Equipped with this technical advance, we observed a facilitation of simultaneous target disappearances and reappearances, strongly implying long-range interactions between the distant targets.

The multi-target display also allowed us to have a more objective graded measure of differences in the contents of consciousness (i.e., the amount of PFI), which revealed an asymmetry between the neural correlates of disappearances and reappearances: background SNR increased with the amount disappearance, but not reappearance during multi-target PFI. One possible explanation is the difference in saliency between PFI disappearances and reappearances, as reappearances can be predicted with higher spatial and temporal accuracy than disappearances. Increased spatial accuracy follows from the fact that reappearances can only occur at locations where a target has already disappeared moments prior. As the duration of PFI is also short compared to the 60-second trial (Figure 7), reappearances can also be predicted with greater temporal accuracy than multi-target disappearances. Thus, PFI disappearances may be more unexpected than reappearances, enhancing their subjective saliency. Indeed, greater phasic pupil responses to target disappearances than reappearances have been reported in motion-induced blindness (Kloosterman et al., 2015; Thomas et al., 2017) which may be closely related to PFI (Devyatko, Appelbaum, & Mitroff, 2017; Hsu, Yeh, & Kramer, 2004, 2006; New & Scholl, 2008). This difference in spatiotemporal saliency might have resulted in the asymmetric patterns of log(SNR) based on the number of disappearing or reappearing targets (Figure 8). To better understand the mechanisms of this asymmetry, further studies employing a paradigm that features multi-target and graded conscious perception will be necessary. Another possibility, unexplored in the present dataset, is the contribution of eye-movements to PFI phenomena. Future studies could address this limitation and test whether the pattern of saccades and microsaccades differ when targets are visible or invisible during PFI, and whether these potential differences affect visibility and/or the neural correlates measured as SSVEPs.

### Insights into PFI mechanisms

Our results are relevant to two popular models of PFI. The first is an isomorphic model. This model proposes the primary substrate of PFI are neurons in early retinotopic areas corresponding to target regions, which are activated via neurons corresponding to their surrounds through lateral connections (De Weerd et al., 1995; Pessoa et al., 1998). The model specifically proposes a two-stage process, where a first stage of seconds-long boundary adaptation is followed by a second stage of near instantaneous interpolation of the target location by surrounding visual features (Spillmann & De Weerd, 2003). The second is a symbolic model, whereby filling-in occurs when the visual system ignores an absence of information (Dennett, 1991; Kingdom & Moulden, 1988; O’ Regan, 1992). In this model, the phenomenon of filling-in is realized at a (possibly higher) representational level, whereby a region devoid of information is symbolically labelled as ‘more of the same’ background, and thus is rendered invisible.

In favour of the isomorphic model, previous electrophysiological data has recorded increased spike rates in regions responding to a filled-in pattern in monkeys (De Weerd et al., 1995). Importantly, De Weerd et al.’s (1995) single-unit study did not obtain subjective reports from the monkeys. With human participants, we provide this critical aspect of the data on an event-by-event manner. This allowed us to examine if the exact timing of the increases in neural activity precedes or follows the onset of a perceptual disappearance. By recording simultaneous subjective reports, we conclude that an increase in background SNR precedes PFI events. This slow, seconds-long increase in background-related SNR prior to PFI events supports an active mechanism as a catalyst for PFI, which is central to the isomorphic model.

On the other hand, the symbolic model that suggests that filling-in happens in higher-level visual areas (Pessoa et al., 1998) is also consistent with our behavioural findings. We observed a synergistic effect among spatially distant targets, which implies the involvement of neurons that have larger receptive fields, typically found only in higher-level visual areas (Dumoulin & Wandell, 2008; Yoshor, Bosking, Ghose, & Maunsell, 2007). This across-quadrant faciliatory interaction extends a previous report of within-quadrant interactions during PFI (De Weerd et al., 1998, experiment 4). More specifically, this synergistic PFI across quadrants may point to a mechanism that facilitates perceptual grouping (Wagemans et al., 2012).

Grouping may also interact with attentional mechanisms. Indeed, attending to shared features such as temporal modulation has been shown to enhance the binding of distributed visual regions into a perceptual group (Alais, Blake, & Lee, 1998). As attending to shared features such as colour (Lou, 1999) or shape (De Weerd et al., 2006) increases the disappearance of peripherally presented targets, fluctuations in attention to the targets as a group may also have impacted on multiple-locations synergistically. Alternatively, the simultaneous disappearance of multiple targets could be due to random fluctuations of the brain’s response to the background (potentially also modulated by attention). Since the background surrounds all targets, a temporary increase in response could affect the visibility of all targets simultaneously.

Overall, our results are not compatible with the view that PFI is a phenomenon that results purely due to local adaptation processes in the retinal or low-level visual areas. Instead our results are compatible with the view that both retinotopic and contextual influences, possibly through lateral connections, determine the dynamics of PFI (Sasaki, 2007).

### Spatiotemporal profiles of 1f and 2f background SSVEP are distinct

Another insight that arose from our application of SSVEP to study PFI regards the difference in spatiotemporal profiles of 1f and 2f responses (Figure 10 and 11). This difference was specifically modulated around the time of PFI. In the literature, 1f and 2f are traditionally considered to be similar, as they are dictated by the same stimulus input (Norcia et al., 2015). Recently, this assumption has been challenged by the finding of an attentional modulation of 2f, but not 1f, with concomitant changes in hemispheric lateralization for the topography of SSVEP responses (Kim et al., 2011; Kim & Verghese, 2012). While an increased spatial distribution of 2f compared to 1f is consistent with our results where 1f was strongest over mid-occipital sites and 2f extended laterally (Figure 5), the flicker stimuli used in our experiments differ from those studies that optimized differentiating 1f from 2f (Kim et al., 2011). As such, extending this interpretation to our findings should be done with caution, but the temporal advantage of the 2f crossover compared to the 1f crossover would be consistent with a covert attentional modulation of 2f that instigates a perceptual change. Future studies with an explicit attentional manipulation will be needed to confirm whether the harmonic differences we have reported are due to the allocation of attention.

### Conclusions

Here we extend efforts to refine NCC paradigms, by using PFI. Unlike traditional stimuli, PFI has the advantage that perceptual changes can be easily mimicked physically, and that participants can accurately report on multiple changes in consciousness occurring in close temporal proximity without much training. While genuine PFI and physical PMD periods were phenomenally similar, we revealed significant differences in their respective neural substrates through our SNR reconstruction analysis, and suggest that these differences are due to the presence of competitive mechanisms supporting perceptual disappearances, but not physical disappearances. Future studies that succeed in tagging both targets and surrounds in PFI would be able to investigate the nature of this competition. They may also reveal why there are significant differences in the dependence on the amount of PFI for disappearances, but not reappearances, which we have tentatively linked to differences in the level of expectation and saliency. These are intriguing empirical questions to be resolved in the future by capitalizing upon the peculiar effect that attention increases PFI (De Weerd et al., 2006; Lou, 1999) and/or by utilizing SSVEP-based no-report paradigms (Tsuchiya et al., 2015). We hope that our approach that combines under-utilized PFI with SSVEP techniques will inspire various novel designs to address this central question in cognitive neuroscience: the neural basis of attention and consciousness.

## Acknowledgements

MJD was supported by an Australian Government Research Training Program Scholarship and by an Australian Research Council (ARC) Discovery Project (DP) (DP180104128). NT was funded by an ARC Future Fellowship (FT120100619) and DPs (DP130100194, DP180100396, and DP180104128).

## Notes

**Conflict of Interest:** The authors declare no competing financial interests.

#### Summary of Updates

This version of the manuscript has been revised to include new analyses of SNR timing differences, SNR activity during PMD periods, clarification of our novel SNR reconstruction procedure, and clarification of figure legends.

